# Characterization of the *Brassica napus* flavonol synthase gene family reveals bifunctional flavonol synthases

**DOI:** 10.1101/2021.06.30.450533

**Authors:** Hanna Marie Schilbert, Maximilian Schöne, Thomas Baier, Mareike Busche, Prisca Viehöver, Bernd Weisshaar, Daniela Holtgräwe

**Affiliations:** Genetics and Genomics of Plants, CeBiTec & Faculty of Biology, Bielefeld University, 33615 Bielefeld, Germany; Algaebiotechnology and Bioenergy, CeBiTec & Faculty of Biology, Bielefeld University, 33615 Bielefeld, Germany

**Keywords:** flavonoid biosynthesis, specialized metabolism, rapeseed, 2-oxoglutarate-dependent dioxygenases, flavanone 3-hydroxylase, bifunctionality, gene family

## Abstract

Flavonol synthase (FLS) is a key enzyme for the formation of flavonols, which are a subclass of the flavonoids. FLS catalyses the conversion of dihydroflavonols to flavonols. The enzyme belongs to the 2-oxoglutarate-dependent dioxygenases (2-ODD) superfamily. We characterized the *FLS* gene family of *Brassica napus* that covers 13 genes, based on the genome sequence of the *B. napus* cultivar Express 617. The goal was to unravel which *BnaFLS* genes are relevant for seed flavonol accumulation in the amphidiploid species *B. napus*. Two *BnaFLS1* homeologs were identified and shown to encode bifunctional enzymes. Both exhibit FLS activity as well as flavanone 3-hydroxylase (F3H) activity, which was demonstrated *in vivo* and *in planta. BnaFLS1-1* and *-2* are capable of converting flavanones into dihydroflavonols and further into flavonols. Analysis of spatio-temporal transcription patterns revealed similar expression profiles of *BnaFLS1* genes. Both are mainly expressed in reproductive organs and co-expressed with the genes encoding early steps of flavonoid biosynthesis. Our results provide novel insights into flavonol biosynthesis in *B. napus* and contribute information for breeding targets with the aim to modify the flavonol content in rapeseed.

## 1 Introduction

Rapeseed (*Brassica napus* L.) is the second most important oil crop worldwide (Nesi et al., 2008; OECD-FAO and Connell, 2015). The high oil (∼50%) and protein (∼25%) content of *B. napus* seed is the result of decades of extensive breeding aiming to improve its nutritional quality and agronomical yield (Nesi et al., 2008). Still, the presence of anti-nutritional components, like phenolic compounds or glucosinolates, render rapeseed protein essentially unusable for human consumption (Wang et al., 2018; Hald et al., 2019). While glucosinolate break-down products cause metabolic disturbances, phenolics can impair digestibility and cause a strong bitter off-taste (Nesi et al., 2008; Wanasundara et al., 2016; Hald et al., 2019). The glucosinolates amount in seeds have been greatly reduced through breeding of double zero lines with improved nutraceutical properties (Nesi et al., 2008). However, breeding of low phenolic lines with optimal compositions for the use of rapeseed protein as edible vegetable product is difficult. The reason is the great diversity of phenolic compounds and their involvement in many processes which impact plant fitness (Auger et al., 2010; Wang et al., 2018). Phenolics can be beneficial for human health due to their antioxidant activity, thereby facilitating the prevention of cardiovascular diseases and cancer (Wang et al., 2018). On the other hand, phenolics can i) impair digestibility, ii) cause undesired dark color, and iii) cause bitter off-taste derived from kaempferol-derivatives (Auger et al., 2010; Hald et al., 2019). Therefore, breeding of low or high phenolic cultivars depends on their economic use, e.g. use as seed oil/animal feed or edible vegetable (Wang et al., 2018).

Flavonoids are a major group of phenolics and belong to a diverse class of plant specialized metabolites comprising over 9,000 different substances (Williams and Grayer, 2004; Grotewold, 2006). They are derived from flavonoid biosynthesis (Figure 1), which branch of from the phenylalanine-based general phenylpropanoid pathway (Hahlbrock and Scheel, 1989). Flavonoids are classified in different subgroups, namely chalcones, flavones, flavandiols, anthocyanins, proanthocyanidins (PAs), aurones, and flavonols (Winkel-Shirley, 2001). Flavonols define the largest subgroup of flavonoids, mainly due to a plethora of glycosylation patterns (Zhang et al., 2013). They are classified in e.g. kaempferols and quercetins depending on the hydroxylation pattern of the B ring (Winkel-Shirley, 2001). Flavonols are colorless for the human eye but absorb in the ultraviolet (UV) range. After light treatment, they accumulate in their glycosylated form in the vacuole of epidermal and mesophyll cells or on occasion in epicuticular waxes (Weisshaar and Jenkins, 1998; Winkel-Shirley, 2001; Agati et al., 2009). Their biosynthesis is largely influenced by environmental cues such as temperature and UV light (Winkel-Shirley, 2002; Olsen et al., 2009). Flavonols have several physiological functions in plants including antimicrobial properties, UV protection, modulation of auxin transport, male fertility, and flower pigmentation together with anthocyanins (Harborne and Williams, 2000; Peer and Murphy, 2007).

**Figure 1:**
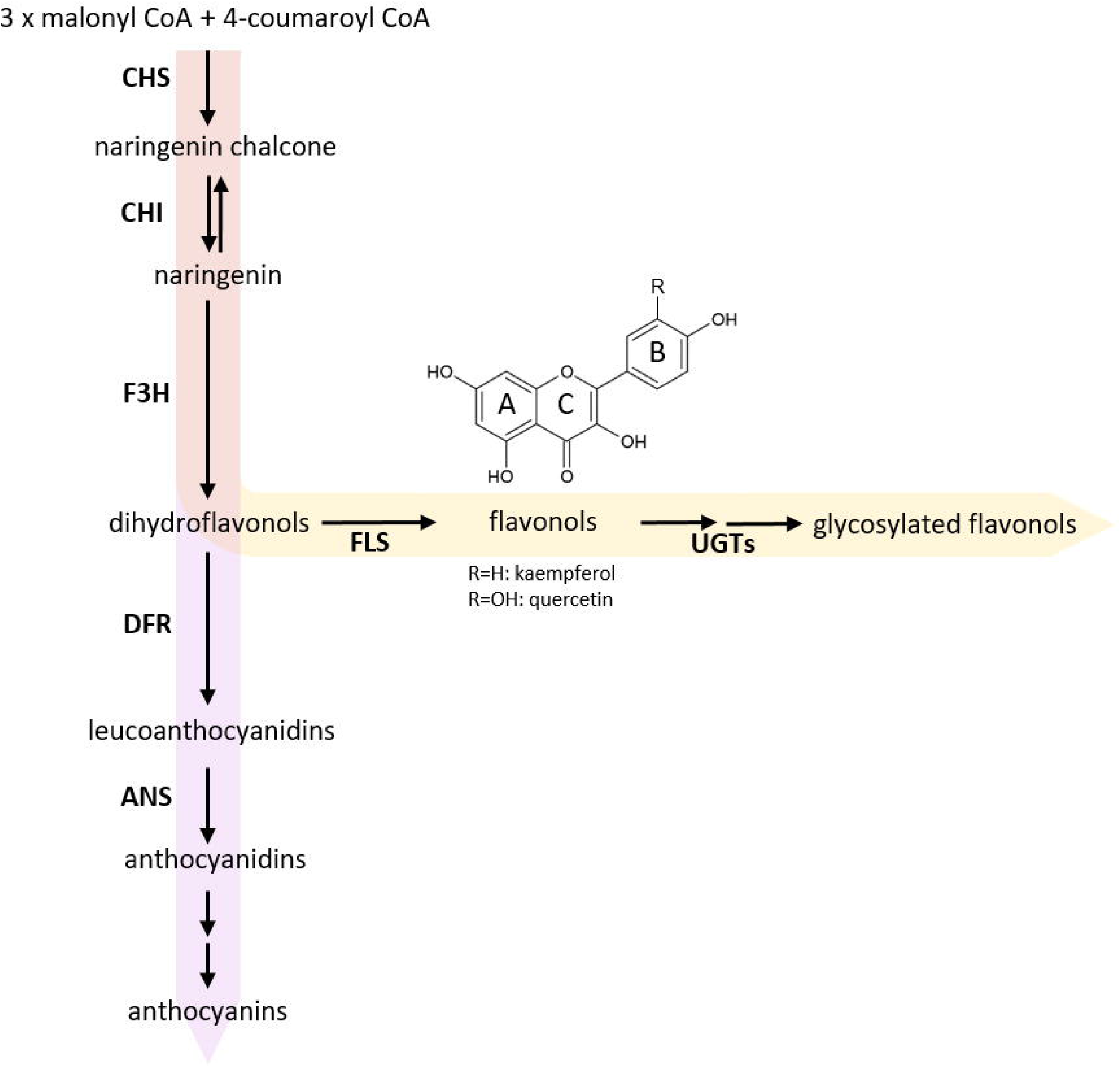
Simplified scheme of flavonoid biosynthesis. The flavonol biosynthesis pathway (highlighted via an orange arrow) is part of the flavonoid biosynthesis, which also includes the anthocyanin pathway (highlighted via a violet arrow) (modified after (Winkel-Shirley, 2001)). The metabolic flux into the flavonol biosynthesis is influenced by dihydroflavonol 4-reductase (DFR) as it competes with FLS for substrates. Enzyme names are abbreviated as follows: chalcone synthase (CHS), Chalcone isomerase (CHI), flavanone 3-hydroxylase (F3H), flavonol synthase (FLS), UDP-glycosyltransferases (UGTs), anthocyanidin synthase (ANS).

The central enzyme of flavonol biosynthesis is flavonol synthase (FLS). FLS converts a dihydroflavonol into the corresponding flavonol by introducing a double bond between C-2 and C-3 of the C-ring (Figure 1)(Forkmann et al., 1986; Holton et al., 1993). FLS activity was first identified in irradiated parsley cells (Britsch et al., 1981). Several studies identified more than one *FLS* gene in the genome of a given species, including *Zea mays* (Falcone Ferreyra et al., 2012), *Musa acuminata* (Busche et al., 2021), *Vitis vinifera* (Downey et al., 2003; Fujita et al., 2006), *Fressica hybrida* (Shan et al., 2020), and *Arabidopsis thaliana* (Pelletier et al., 1997; Owens et al., 2008). In *A. thaliana*, which is evolutionary closely related to *B. napus*, most genes of the central enzymes of the flavonoid biosynthesis are encoded by single-copy genes. However, *FLS* marks an exception as there are six genes annotated in the *A. thaliana* genome sequence (Pelletier et al., 1997; Owens et al., 2008). Only *FLS1* encodes a functional FLS, thus being the major contributor to flavonol production in *A. thaliana* (Wisman et al., 1998). It has been postulated that the *AthFLS* gene family derived from recent gene duplication events and is currently undergoing a pseudogenisation process to eliminate ‘unnecessary’ gene copies (Preuss et al., 2009; Stracke et al., 2009). The Brassicaceae-lineage specific whole genome triplication followed by diploidization after divergence from the common ancestor of *A. thaliana* and *B. napus* (Wang et al., 2011; Chalhoub et al., 2014) suggests that the amphidiploid *B. napus* harbours an even larger *FLS* family, which formally may cover up to 36 members. So far, six *FLS* genes have been identified for the A-subgenome donor *B. rapa* (Guo et al., 2014), while the C-subgenome donor *B. oleracea* has not yet been studied in detail. Up to now, the exact size of the *B. napus FLS* gene family remains unknown. Previous studies on the flavonol biosynthesis in *B. napus* were mainly focused on metabolites (Auger et al., 2010) or covered transcriptomic and phylogenetic analysis of genes preceding the FLS reaction in the flavonol pathway (Qu et al., 2016).

Some FLSs have been characterized as bifunctional enzymes, exhibiting FLS and F3H activity (Figure 1), e.g. in *A. thaliana* (Prescott et al., 2002; Owens et al., 2008), *Oriza sativa* (Park et al., 2019), *Citrus unshiu* (Lukacin et al., 2003), and *Ginkgo biloba* (Xu et al., 2012). FLS has been classified as a 2-oxoglutarate-dependent dioxygenase (2-ODD), similar to flavanone 3-hydroxylase (F3H) and anthocyanidin synthase (ANS). The three enzymes display partial amino acid (aa) sequence similarity and overlapping functions (Prescott and John, 1996; Cheng et al., 2014). The nonheme cytosolic 2-ODD enzymes require 2-oxoglutarate as co-substrate, while ferrous iron acts as co-factor (Cheng et al., 2014). FLS and ANS are relatively closely related with 50-60% aa sequence similarity, while F3H share less than 35% similarity with FLS and ANS (Lukacin et al., 2003; Cheng et al., 2014). ANS, an enzyme catalyzing a late step in the flavonoid biosynthesis pathway (Figure 1), can have both FLS and F3H activity (Welford et al., 2001; Cheng et al., 2014). Therefore, ANS contributes to flavonol production, although (at least in *A. thaliana*) to a much lesser extent than FLS (Preuss et al., 2009). In addition, 2-ODDs display species-specific substrate specificities and affinities (Preuss et al., 2009; Park et al., 2017; Jiang et al., 2020).

The transcriptional regulation of flavonol biosynthesis is mainly achieved by the combinatorial action(s) of MYB11, MYB12, and MYB111, which belong to subgroup 7 (SG7) of the R2R3-MYB transcription factor family (Mehrtens et al., 2005; Stracke et al., 2007). However, the *myb11/myb12/myb111* triple mutant of *A. thaliana* retains its pollen flavonol composition (Stracke et al., 2010). This led to the discovery of MYB99, MYB21, and MYB24, which together control flavonol biosynthesis in anthers and pollen (Battat et al., 2019; Shan et al., 2020). MYB21, MYB24, and the SG7 MYBs function as independent transcriptional activators (Mehrtens et al., 2005; Stracke et al., 2007; Shan et al., 2020). The SG7 MYBs can activate all genes belonging to flavonol biosynthesis including *CHS, CHI, F3H*, and *FLS* (Mehrtens et al., 2005; Stracke et al., 2007). Recently, direct activation of *AthFLS1* by AthMYB21 and AthMYB24 was shown in *A. thaliana* (Shan et al., 2020).

In this study, we characterize 13 members of the *BnaFLS* gene family, which is one of the largest FLS enzyme families analyzed to date. We separated the *BnaFLS* genes from *F3H* and *ANS* genes of *B. napus*. Only one *FLS* gene has been characterized so far in *B. napus* (Vu et al., 2015). We demonstrate that both *BnaFLS1* homeologs encode bifunctional enzymes, exhibiting FLS and F3H activity, while two *BnaFLS3* homeologs encode proteins with solely F3H activity. Moreover, we provide insights into the spatio-temporal transcription of *BnaFLSs* and present hypotheses about the mechanisms underlying FLS bifunctionality. Thus, our study provides novel insights into the flavonol biosynthesis of *B. napus* and supports targeted engineering of flavonol content, e.g. to enable the use of rapeseed protein in human consumption.

## 2 Materials and Methods

### 2.1 Plant material

We used the *B. napus* Express 617, a dark-seeded winter cultivar (Lee et al., 2020). *B. napus* was first grown in the greenhouse under long day conditions and then transferred outside for natural vernalisation, followed by additional growth outside. *A. thaliana* Columbia 0 (Col-0, NASC ID N1092) and Nössen-0 (Nö-0, NASC ID N3081) were used as wildtype controls. The *f3h* mutant (*tt6-2*, GK-292E08, NASC ID N2105575, Col-0 background) (Appelhagen et al., 2014) and the *ans/fls1* double mutant (synonym *ldox/fls1-2, ldox*: SALK_028793, NASC ID N2105579, Col-0 background; *fls1-2*: RIKEN_PST16145, Nö-0 background) (Stracke et al., 2009) were used for the generation of transgenic lines. *A. thaliana* plants were grown in the greenhouse under a 16-h-light/8-h-dark cycle at 22 °C before transformation.

### 2.2 Identification of *BnaFLS* candidate genes

BnaFLS homologs were identified with KIPEs v0.255 as described previously (Pucker et al., 2020). KIPEs was run with a minimal BLAST hit similarity of 40% to reduce the number of fragmented peptides derived from possible mis-annotations. As bait, peptide sequences from the sequence collection of functional F3H, FLS, and ANS sequences described in KIPEs were used. As subject species, the peptide sequence sets of several *Brassica* species were used (Supplementary Table S1). The alignment was constructed with MAFFT v.7 (Katoh and Standley, 2013) and trimmed to minimal alignment column occupancy of 10%. Next, a phylogenetic tree was built with FastTree v2.1.10 (Price et al., 2009) using 10,000 rounds of bootstrapping, including the bait sequences and 2-ODD-like sequences from *A. thaliana* derived from Kawai *et al*. 2014 (Kawai et al., 2014) (Supplementary File S1). The phylogenetic tree was visualized with FigTree v1.4.3 (http://tree.bio.ed.ac.uk/software/figtree/)(Supplementary Figure S1). Classification of BnaFLS candidates was generated based on the corresponding *A. thaliana* orthologs.

### 2.3 Sequence-specific analyses of *BnaFLS* candidates and secondary structure modelling

A comprehensive summary about gene-specific features of *BnaFLS* candidates is summarized in Supplementary Table S2. GSDS 2.0 (Hu et al., 2015) was used to generate gene structure plots. Literature knowledge was used to identify MYB-recognition elements (MRE) within 1 kbp upstream of the translational start site of *BnaFLS* candidates (Supplementary Figure S2). The conserved MRE consensus sequence 5’-AcCTACCa-3’, identified as a SG7 recognition motif (Hartmann et al., 2005; Stracke et al., 2007) and the sequence motifs important for the binding of AthMYB21 (MYBPZM: 5’-CCWACC-3’) and AthMYB24 (MYBCORE: 5’-CNGTTR-3’) to *AthFLS1* were used for screening (Battat et al., 2019; Shan et al., 2020).

Theoretical isoelectric points, as well as molecular weight values of the BnaFLS protein sequences were calculated with ExPASY V (Gasteiger et al., 2005)(Supplementary Table S3). In addition, SignalP v. 5.0 (Almagro Armenteros et al., 2019b) and TargetP v. 2.0 (Almagro Armenteros et al., 2019a) were used to infer the presence of signal peptides and N-terminal presequences of BnaFLS candidates, respectively (Supplementary Table S4, S5). TMHMM v. 2.0 (Krogh et al., 2001) was used to predict transmembrane regions within BnaFLS sequences (Supplementary Table S2). Finally, Plant-mPLoc v. 2.0 (Chou and Shen, 2010) was used to predict the subcellular localization of BnaFLS candidates (Supplementary Table S2). Amino acid sequence identities of BnaFLSs compared to FLS homologs of *A. thaliana, B. rapa*, and *B. oleracea* were calculated based on a MAFFT alignment (Supplementary Table S6; https://github.com/hschilbert/BnaFLS). Protein sequence alignments were visualised at http://espript.ibcp.fr/ESPript/ESPript/index.php v. 3.0 (Robert and Gouet, 2014) using the AthFLS1 pdb file derived from Pucker *et al*. 2020 (Pucker et al., 2020). Functionally relevant amino acid residues and motifs for FLS and F3H activity were highlighted.

*In silico* secondary structure models of relevant BnaFLS candidates were generated via I-TASSER (Roy et al., 2010) and visualized with Chimera v. 1.13.1 (Pettersen et al., 2004). The AthF3H PDB file derived from Pucker *et al*. 2020 (Pucker et al., 2020) was used for visualisation. The generated PDB files of this work can be accessed via Supplementary File S2.

### 2.4 Gene expression analysis: Ribonucleic acid extraction, library construction, and sequencing

Ribonucleic acid (RNA) samples were isolated from seeds and leaves using the NucleoSpin^®^ RNA Plant kit (Macherey-Nagel, Düren, Germany) according to manufacturer’s instructions. Seed samples of the *B. napus* cultivar Express 617 were collected 23 and 35 days after flowering (DAF), while leave samples were collected 35 DAF. Samples were collected in triplicates. The RNA quality was validated using NanoDrop and Agilent 2100 to confirm the purity, concentration, and integrity, respectively. Based on 1 μg of total RNA, sequencing libraries were constructed following the TruSeq v2 protocol. Three seed and leaf samples per genotype were processed. Single end sequencing of 82 nt was performed on an Illumina NextSeq 500 at the Sequencing Core Facility of the Center for Biotechnology (CeBiTec) at Bielefeld University.

### 2.5 Gene expression analysis and co-expression analysis using *B. napus* RNA-Seq data

Read quality was assessed by FastQC (Andrews, 2018), revealing reads of good quality reaching a phred score of 35 or above. Next, reads were mapped to the Express 617 reference genome sequence (Lee et al., 2020) using STAR v. 2.7.1a (Dobin et al., 2013). STAR was run in basic mode allowing maximal 5% mismatches per read length and using a minimum of 90% matches per read length. These read mappings were used to manually correct the functional annotation of the *BnaFLS* candidates (Supplementary File S3). The corresponding corrected annotation file was used for downstream analysis.

Beside the newly generated RNA-Seq data, publicly available RNA-Seq data sets were used and retrieved from the Sequence Read Archive (https://www.ncbi.nlm.nih.gov/sra) via fastq-dump v. 2.9.6 (https://github.com/ncbi/sra-tools) to analyze the expression of the candidate genes across various organs (Supplementary Table S7). Kallisto v. 0.44 (Bray et al., 2016) was used with default parameters to quantify transcripts abundance. The heatmap was constructed with a customized python script (https://github.com/hschilbert/BnaFLS) using mean transcripts per millions (TPMs) per organ. Condition-independent co-expression analysis was performed to identify co-expressed genes using Spearman’s correlation coefficient (https://github.com/hschilbert/BnaFLS) by incorporating 696 RNA-Seq data sets (Supplementary Table S8). To filter for strong co-expression the Spearman’s correlation coefficient threshold was set to 0.7 as suggested by Usadel *et al*. 2009 (Usadel et al., 2009).

### 2.6 Functional annotation of *B. napus* Express 617 genes

Genes were functionally annotated by transferring the *A. thaliana* Araport11 (Cheng et al., 2017) functional annotation to the *B. napus* Express 617 gene models. The annotation was used for the co-expression analysis. OrthoFinder v. 2.3.7 (Emms and Kelly, 2019) was applied using default parameters to identify orthogroups between the representative peptide sequences of Araport11 and the *B. napus* Express 617 peptide sequences as previously defined (Pucker et al., 2017). Remaining nonannotated genes were functionally annotated by using reciprocal best blast hits (RBHs) and best blast hits (BBHs) as described previously (Pucker et al., 2016)(Supplementary Table S9).

### 2.7 Generation of *BnaFLSs* constructs

All constructs generated in this work were produced via Gateway cloning technique according to manufacturer’s instructions and verified by DNA sequencing (Supplementary Table S10). Total RNA from leaves and seeds of Express 617 was extracted as described above (see 2.4). Complementary DNA (cDNA) was synthesized with the ProtoScript™ Reverse Transcriptase kit (Invitrogen, Karlsruhe, Germany) using ∼1 μg of total RNA and 1 μl of oligo (dT) and 1 μl of random-hexamer primers. cDNA fragments corresponding to the full-length ORFs of the candidate genes were then amplified via PCR with Q5^®^ High-Fidelity Polymerase PCR kit (NEB, Frankfurt am Main, Germany) using gene-specific gateway primers (Supplementary Table S9). The sizes of the amplification products were analyzed by gel electrophoresis and visualized by ethidium bromide on a 1% agarose gel. The amplicons were purified from the PCR reagent tube via the NucleoSpin^®^ Gel and PCR Clean-up Kit (Macherey-Nagel, Düren, Germany).

The purified cDNA fragments corresponding to the full-length ORFs of the candidate genes were then recombined into *pDONR*™*/Zeo* (Invitrogen, Karlsruhe, Germany) using the Gateway BP Clonase II Enzyme Mix (Invitrogen, Karlsruhe, Germany) and the *attB* recombination sites of the respective gateway primers (Supplementary Table S10). Each entry clone was then used to transfer the CDS into the destination vector *pLEELA* (Jakoby et al., 2004) or *pDEST17* (Invitrogen) via the Gateway LR Clonase II Enzyme Mix (Invitrogen, Karlsruhe, Germany). In *pLEELA*, the rapeseed coding sequences are under control of a double 35S promoter. *pDEST17* was used for heterologous protein expression during the *in vivo E. coli* bioconversion assay under the control of the T7 promotor. The following constructs were available from previous studies: *pDEST17-AthF3H, pDEST17-AthFLS1* (Busche et al., 2021), *pDONR-AthFLS3, pDONR-AthFLS5, pDONR-AthANS* (Preuss et al., 2009). The respective *BnaFLS* CDS sequences are listed in Supplementary File S4.

### 2.8 F3H and FLS bioconversion assay in *E. coli*

The bioconversion assay in *E. coli* subsequent HPTLC analysis of the methanolic extracts were performed as described in Busche *et al*. 2021 (Busche et al., 2021). Successful heterologous expression of the recombinant proteins via SDS-PAGE was shown (Supplementary Figure S3).

### 2.9 Generation of complementation lines

The generated *pLEELA-BnaFLSX* constructs were used to transform the *A. thaliana f3h* knock out mutant, as well as the *ans*/*fls1* double mutant using the *A. tumefaciens* strain GV3101::pM90RK (Koncz and Schell, 1986) according to the floral dip protocol (Clough and Bent, 1998). Selection of T1 plants was carried out by BASTA selection. Surviving plants were genotyped for the respective wildtype and mutant alleles, as well as the insertion of the transgene into the genome and its expression via PCR and RT-PCR (Supplementary Table S10). The genotyping for the presence of the transgene was repeated with T2 plants. T2 plants were used for the generation of flavonol-containing methanolic extracts as described below (see 2.10). T2 plants of the transformed *ans/fls1* mutants and T3 plants of the transformed *f3h* mutants were used for DPBA-staining of young seedlings (see 2.11).

### 2.10 Flavonol content analysis by high-performance thin-layer chromatography (HPTLC)

The flavonol glycosides were extracted and analyzed as previously described (Stracke et al., 2009). *A. thaliana* stems were homogenised in 80% methanol and incubated for 15 min at 70 °C and then centrifuged for 10 min at 16,100 xg. The supernatants were vacuum-dried at 60 °C and sediments were dissolved in 1 μl of 80% methanol mg^-1^ starting material for HPTLC analysis. In total, 3 μl of each sample were spotted on silica-60 HPTLC-plates. The *A. thaliana* accessions Col-0 and Nössen-0, as well as the *ans*/*fls1* double mutant were used as controls for the *ans*/*fls1 A. thaliana* complementation lines. For the *f3h* complementation lines, Col-0 and the *f3h* mutant were used as controls. The mobile phase consisted of a mixture of 66.7% ethyl acetate, 8% formic acid, 8% acetic acid, and 17.3% water. Flavonoid compounds were detected as described before (Stracke et al., 2009).

### 2.11 *In situ* flavonoid staining of whole seedlings

The visualisation of flavonoids via DPBA-staining with whole seedlings was performed as described (Stracke et al., 2007), with the following minor adaptations: the bleached seedlings were stained to saturation in a freshly prepared aqueous solution of 0.25% (w/v) DPBA, 0.01% (v/v) Triton X-100, and 20% ethanol (v/v).

## 3 Results

### 3.1 FLS family of *B. napus*

We identified a monophyletic group of 13 BnaFLS candidates through phylogenetic analysis using F3H, ANS, and 2-ODD-like protein sequences as outgroup to classify members of the 2-ODD family (Figure 2, Supplementary Figure S1, Supplementary Table S2) of *B. napus*. The BnaFLS candidates were further classified within the *FLS* gene family based on their phylogenetic relationship to their most likely *A. thaliana* orthologs (Figure 2). Thereby, we identified two BnaFLS1, two BnaFLS2, five BnaFLS3, and four BnaFLS4 candidates in the *B. napus* cultivar Express 617. *BnaFLS1-1* was identified on chromosome C09, while its homeolog *BnaFLS1-2* is located on chromosome A09.

**Figure 2:**
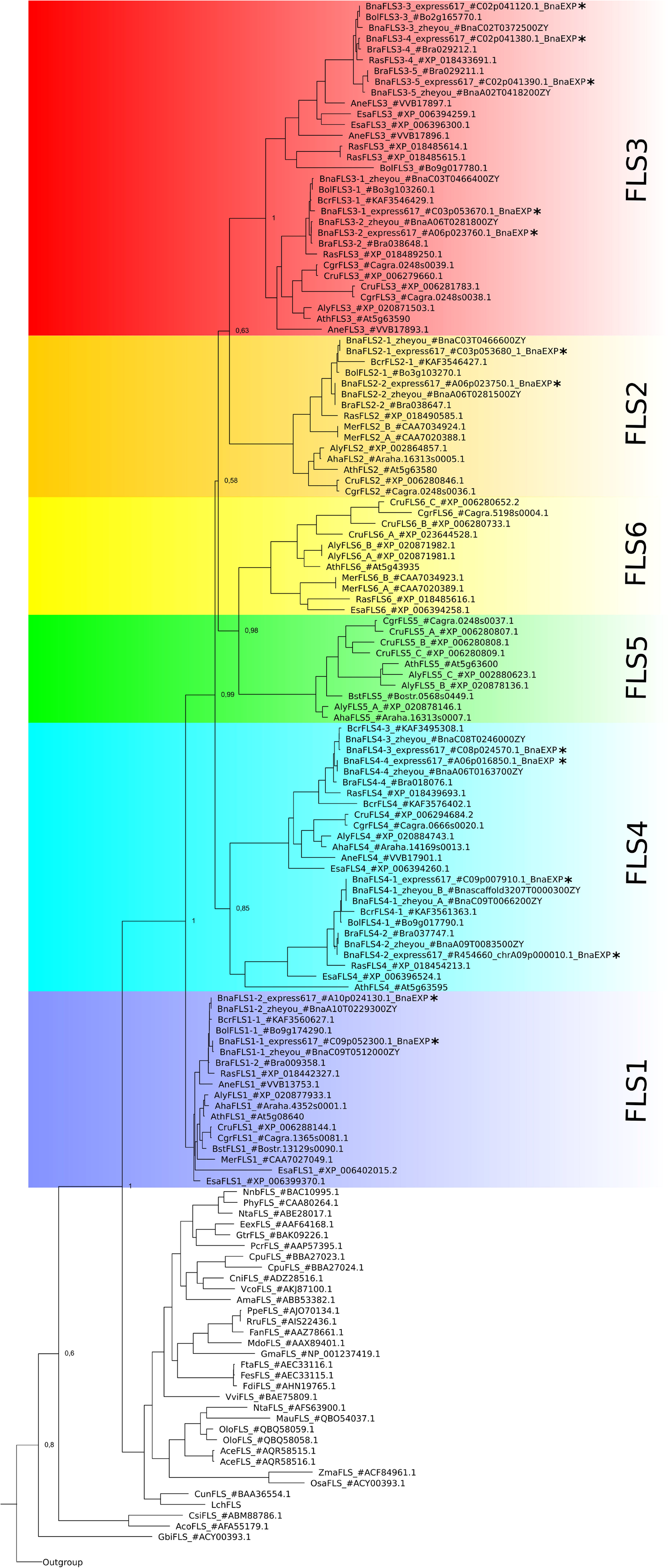
Phylogeny of BnaFLS candidates and previously described FLS sequences. Relative bootstrap-values are shown next to relevant nodes. The phylogenetic tree is based on amino acid sequences. FLS family members of *B. napus* Express 617 are marked with an asterisk. The outgroup comprises the 2-ODD members ANS and F3H, as well as 2-ODD-like sequences (Supplementary Figure S1).

The genomic structure of the *BnaFLS* candidate genes comprises 3-4 exons and the encoded proteins display a length range from 270 to 336 amino acids (aa) (Table 1, Supplementary Figure S4, Supplementary Table S2). Considering the chromosomal rearrangements as described for the cultivar Darmor-bzh (Chalhoub et al., 2014), homeologs were identified (Table 1).

**Table 1:** Chromosomal location of *BnaFLS* candidate genes in Express 617. The genomic position and exon number per *BnaFLS* candidate gene based on the *B. napus* Express 617 assembly are listed. Moreover, the amino acid (AA) length of the corresponding protein is stated. Homeologs are located inside one row.

| Gene name | Chromosome | Position [kbp] | No. of exons | AA length |
| --- | --- | --- | --- | --- |
| *BnaFLS1-1* | C09 | 57,490 - 57,492 | 3 | 336 |
| *BnaFLS1-2* | A10 | 18,238 - 18,240 | 3 | 336 |
| *BnaFLS2-1* | C03 | 45,458 - 45,461 | 3 | 307 |
| *BnaFLS2-2* | A06 | 21,674 - 21,677 | 3 | 307 |
| *BnaFLS3-1* | C03 | 45,437 - 45,438 | 4 | 270 |
| *BnaFLS3-2* | A06 | 21,693 - 21,694 | 3 | 297 |
| *BnaFLS3-3* | C02 | 49,747 - 49,749 | 3 | 309 |
| *BnaFLS3-4* | C02 | 49,966 - 49,969 | 3 | 309 |
| *BnaFLS3-5* | C02 | 49,972 - 49,974 | 3 | 310 |
| *BnaFLS4-1* | C09 | 5,509 - 5,511 | 3 | 320 |
| *BnaFLS4-2* | A09* | 8 - 11 | 3 | 306 |
| *BnaFLS4-3* | C08 | 33,122 - 33,123 | 3 | 305 |
| *BnaFLS4-4* | A06 | 10,416 - 10,417 | 3 | 305 |

\*unanchored but assigned.

No *FLS5* and *FLS6* homologs were identified in *B. rapa, B. oleracea*, and *B. napus* (Figure 2). Additionally screened *B. napus* cultivars (Gangan, No2127, Quinta, Shenglii, Tapidor, Westar, ZS11, Zheyou7) were in line with these results. As a *FLS6* homolog is present in *Raphanus sativus*, a very close relative to *B. rapa, B. oleracea* and *B. napus*, the latter three might have lost *FLS6* very recently. *FLS5* was not found in the analyzed species of Brassiceae, Arabideae, Eutremeae, and Coluteocarpeae, while at least one copy was present in Camelineae and Boechereae indicating that *FLS5* might have recently emerged in the latter tribes.

### 3.2 Organ- and temporal-specific expression of *BnaFLS* candidates

The expression of all *BnaFLS* candidate genes was analyzed by newly generated and publicly available RNA-Seq data (Table 2, Supplementary Table S7). As seeds are the major organ for agronomical relevance, we screened for *BnaFLS* candidates expressed in seeds. In total, five genes were found to be expressed in seeds: *BnaFLS1-1, BnaFLS1-2, BnaFLS2-1, BnaFLS3-3*, and *BnaFLS3-4*. These five *BnaFLS* candidate genes revealed organ- and seed developmental-specific expression patterns (Table 2).

**Table 2:** Organ-specific expression of *BnaFLS* candidate genes. The mean transcripts per millions (TPMs) for each *BnaFLS* candidate gene per organ is listed. Single-end RNA-Seq data generated in this study derived from leaves (35 DAF) and seeds (23 and 35 DAF) of Express 617 are marked with an asterisk. The remaining organs are based on publicly available paired-end *B. napus* RNA-Seq data sets. The number of analyzed data sets per organ is stated via (n=X). The color gradient from white via light blue to dark blue indicates the expression strength with dark blue symbolizing high expression. Abbreviations: days after flowering (DAF), days after pollination (DAP), shoot apical meristem (SAM).

|  | *BnaFLS1-1* | *BnaFLS1-2* | *BnaFLS2-1* | *BnaFLS2-2* | *BnaFLS3-1* | *BnaFLS3-2* | *BnaFLS3-3* | *BnaFLS3-4* | *BnaFLS3-5* | *BnaFLS4-1* | *BnaFLS4-2* | *BnaFLS4-3* | *BnaFLS4-4* |
| --- | --- | --- | --- | --- | --- | --- | --- | --- | --- | --- | --- | --- | --- |
| SAM (n=16) | 2 | 3 | 0 | 0 | 0 | 1 | 27 | 0 | 0 | 1 | 0 | 2 | 1 |
| anther prophase 1 (n=12) | 14 | 3 | 0 | 0 | 0 | 0 | 1 | 3 | 0 | 0 | 0 | 0 | 0 |
| anther bolting (n=6) | 227 | 248 | 2 | 0 | 0 | 0 | 6 | 0 | 0 | 0 | 1 | 6 | 2 |
| anther flowering (n=4) | 148 | 131 | 0 | 0 | 3 | 0 | 13 | 4 | 0 | 0 | 0 | 4 | 1 |
| stamen (n=1) | 6 | 7 | 0 | 0 | 0 | 12 | 12 | 0 | 0 | 0 | 0 | 0 | 0 |
| ovule (n=1) | 7 | 1 | 1 | 0 | 0 | 0 | 53 | 3 | 0 | 0 | 1 | 1 | 0 |
| pistil (n=3) | 20 | 33 | 1 | 1 | 0 | 2 | 16 | 0 | 0 | 0 | 0 | 1 | 0 |
| sepal (n=1) | 12 | 10 | 0 | 0 | 0 | 0 | 5 | 0 | 0 | 0 | 3 | 8 | 5 |
| petal (n=2) | 132 | 150 | 0 | 0 | 0 | 1 | 8 | 2 | 0 | 0 | 0 | 0 | 0 |
| silique 10-20DAF (n=13) | 6 | 10 | 6 | 0 | 0 | 1 | 22 | 6 | 0 | 0 | 0 | 2 | 1 |
| silique 25DAF (n=6) | 7 | 8 | 5 | 1 | 0 | 0 | 29 | 43 | 0 | 0 | 0 | 1 | 0 |
| silique 30DAF (n=6) | 21 | 20 | 3 | 1 | 0 | 0 | 16 | 29 | 0 | 0 | 0 | 0 | 0 |
| silique 40DAF (n=2) | 20 | 57 | 1 | 0 | 0 | 0 | 3 | 10 | 0 | 0 | 0 | 0 | 0 |
| seed 23DAF* (n=3) | 15 | 4 | 1 | 0 | 0 | 0 | 55 | 76 | 0 | 0 | 0 | 1 | 1 |
| seed 35DAF* (n=3) | 59 | 29 | 0 | 0 | 0 | 0 | 14 | 29 | 0 | 0 | 0 | 0 | 0 |
| seed coat 14DAF (n=7) | 14 | 20 | 4 | 0 | 0 | 0 | 82 | 8 | 0 | 1 | 0 | 1 | 0 |
| seed coat 21DAF (n=6) | 39 | 32 | 12 | 0 | 0 | 0 | 53 | 165 | 1 | 0 | 0 | 3 | 1 |
| seed coat 28DAF (n=6) | 25 | 20 | 10 | 0 | 0 | 0 | 11 | 146 | 0 | 1 | 0 | 2 | 1 |
| seed coat 35DAF (n=6) | 15 | 10 | 11 | 0 | 0 | 0 | 14 | 210 | 0 | 3 | 0 | 3 | 1 |
| seed coat 42DAF (n=6) | 10 | 3 | 16 | 0 | 1 | 0 | 24 | 134 | 0 | 4 | 0 | 2 | 1 |
| embryo (n=6) | 7 | 60 | 0 | 0 | 0 | 0 | 12 | 2 | 0 | 0 | 0 | 0 | 0 |
| endosperm (n=8) | 7 | 3 | 1 | 0 | 0 | 1 | 8 | 66 | 0 | 3 | 2 | 0 | 0 |
| seedling (n=9) | 5 | 6 | 1 | 0 | 1 | 9 | 25 | 1 | 0 | 0 | 1 | 11 | 2 |
| leaf 35DAF* (n=3) | 15 | 9 | 0 | 0 | 0 | 0 | 38 | 1 | 0 | 0 | 1 | 0 | 0 |
| stem (n=19) | 9 | 10 | 4 | 4 | 1 | 1 | 40 | 7 | 0 | 0 | 0 | 6 | 2 |
| shoot (n=2) | 6 | 10 | 0 | 0 | 0 | 0 | 23 | 0 | 0 | 0 | 0 | 0 | 0 |
| shoot apices (n=2) | 6 | 5 | 1 | 1 | 0 | 0 | 16 | 12 | 0 | 1 | 0 | 5 | 1 |
| root seedling (n=13) | 0 | 0 | 12 | 3 | 6 | 63 | 36 | 20 | 7 | 2 | 6 | 10 | 4 |
| root 30DAP (n=20) | 0 | 0 | 7 | 5 | 7 | 38 | 22 | 8 | 4 | 1 | 1 | 34 | 31 |
| root 60DAP (n=2) | 0 | 0 | 21 | 10 | 22 | 50 | 107 | 45 | 19 | 1 | 1 | 90 | 59 |

Both *BnaFLS1* candidates revealed similar expression patters, showing the highest expression in late anther development, petals, and seeds. The expression of both *BnaFLS1s* tend to increase in siliques from 10 to 40 days after flowering (DAF). A similar expression pattern was observed in the seed coat revealing a development dependent expression. The biggest differences in *BnaFLS1-1* and *BnaFLS1-2* expression were observed in the embryo, where *BnaFLS1-2* is higher expressed compared to *BnaFLS1-1* indicating organ-specific transcriptional regulation at least for this organ. In contrast to *BnaFLS1s*, both *BnaFLS3s* are only marginally expressed in anthers and petals. While the expression of *BnaFLS1s* peaks during late seed and silique development, the expression of both *BnaFLS3s* peak in the early developmental stages. *BnaFLS3-4* is highly expressed during seed coat development. Contrasting expression patterns of *BnaFLS3-3* and *BnaFLS3-4* were identified in e.g. seed coat samples indicating again organ-specific transcriptional regulation. *BnaFLS2-1* was only marginally expressed in all analyzed organs, showing the highest expression in seed coat and roots. In summary, these findings indicate a role of *BnaFLS1-1, BnaFLS1-2, BnaFLS2-1, BnaFLS3-3*, and *BnaFLS3-4* in seeds.

The five *BnaFLS* candidates expressed in seeds were used for downstream in-depth sequence- and functional analysis of the encoded proteins. The candidates revealed similar genomic structures and an alternative splice variant of *BnaFLS2-1* was detected (Figure 3, Supplementary Table S2, Supplementary Figure S5).

**Figure 3:**
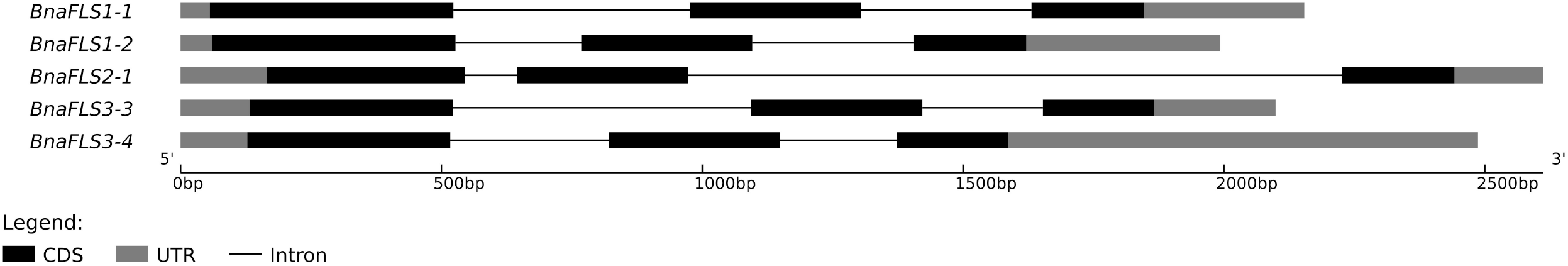
Genomic structure of *BnaFLS* candidates expressed in seeds. The exon-intron structure of *BnaFLS* candidates is shown. The exons are split into coding sequences (CDS, black) and untranslated regions (UTR, gray) and are displayed by rectangles, introns are displayed as black connecting lines.

### 3.3 *BnaFLS1-1* and *BnaFLS1-2* are co-expressed with major players of the flavonoid biosynthesis

To get first insights into which biological pathways the five *BnaFLS* candidates expressed in seeds might be involved, we identified co-expressed genes (Supplementary Table S11, S12, S13, S14, S15). Interestingly, the genes with the most similar expression pattern to *BnaFLS1-1* are part of the flavonoid biosynthesis or the general phenylpropanoid pathway, including *4CL, CHS, CHI, F3H, F3’H, FLS1-2, UGT84A2, GSTF12*, and *MYB111*. Similar results were obtained for *BnaFLS1-2*, which is co-expressed with homolog(s) of *4CL, CHS, CHI, F3H, FLS1-1, UGT84A2*, and *MYB111*. Both *BnaFLS1* genes contain the conserved subgroup 7 MYB-recognition element (MRE) motif in their putative promotor sequences (Supplementary Figure S2).

*BnaFLS3-4* was identified to be co-expressed with genes which mostly lack a functional annotation. However, *BnaFLS3-4* is strongly co-expressed with a *MYB61* homolog. *AthMYB61* is a known regulator of seed coat development. For *BnaFLS2-1* (Spearman’s correlation coefficient < 0.59) and *BnaFLS3-3* (Spearman’s correlation coefficient < 0.69) no genes with strong co-expression could be identified. This is likely due to the very weak expression of *BnaFLS2-1* and the broad expression pattern of *BnaFLS3-3* (Table 2).

### 3.4 BnaFLS candidates share high amino acid sequence identity to *A. thaliana* 2-ODD orthologs

To shed light on the potential functionalities of the *BnaFLS* candidates, the encoded proteins were compared to the well-characterized 2-ODD-members FLS, F3H, and ANS from *A. thaliana* (Table 3). BnaFLS1-1 and BnaFLS1-2 share >91% sequence identity to AthFLS1, while BnaFLS2-1 has 57.4% sequence identity to AthFLS2. BnaFLS3-2 and BnaFLS3-3 revealed a sequence identity of 66.8% to AthFLS3. When comparing all BnaFLS candidates to AthF3H and AthANS, the protein identity ranged from 26.7-31% and 33.6-38.4%, respectively. The two BnaFLS1 candidates share 98.2% sequence identity, differing in 6 aa positions, while both BnaFLS3 candidates have 97.4% sequence identity, differing in 8 aa positions. The high sequence similarity between the BnaFLS candidates and their respective AthFLS orthologs implies close structural relationships and related functions.

**Table 3:** Sequence identity of BnaFLS candidates and 2-ODD members of *A. thaliana*. The protein sequence identity between the BnaFLS candidates and 2-ODD members of *A. thaliana* is given. The heatmap ranging from white via light blue to dark blue indicates low and high sequence identity between the protein pair, respectively. Values are given in percentage.

|  | Ath<br>FLS1 | Ath<br>FLS2 | Ath<br>FLS3 | Ath<br>F3H | Bna<br>FLS1-1 | Bna<br>FLS1-2 | Bna<br>FLS2-1 | Bna<br>FLS3-3 | Bna<br>FLS3-4 | Ath<br>ANS |
| --- | --- | --- | --- | --- | --- | --- | --- | --- | --- | --- |
| AthFLS1 | 100 | 46.4 | 64.1 | 29.8 | 91.1 | 91.4 | 57.7 | 57.3 | 56.7 | 39.9 |
| AthFLS2 |  | 100 | 47.3 | 22.0 | 46.7 | 46.7 | 57.4 | 45.7 | 45.6 | 27.0 |
| AthFLS3 |  |  | 100 | 27.3 | 62.9 | 62.6 | 59.7 | 66.8 | 66.8 | 34.5 |
| AthF3H |  |  |  | 100 | 31.0 | 30.3 | 27.0 | 27.2 | 26.7 | 30.0 |
| BnaFLS1-1 |  |  |  |  | 100 | 98.2 | 56.6 | 56.1 | 55.8 | 38.4 |
| BnaFLS1-2 |  |  |  |  |  | 100 | 56.6 | 55.8 | 55.5 | 38.4 |
| BnaFLS2-1 |  |  |  |  |  |  | 100 | 54.3 | 53.7 | 34.1 |
| BnaFLS3-3 |  |  |  |  |  |  |  | 100 | 97.4 | 33.7 |
| BnaFLS3-4 |  |  |  |  |  |  |  |  | 100 | 33.6 |
| AthANS |  |  |  |  |  |  |  |  |  | 100 |

### 3.5 BnaFLS candidates carry residues important for FLS and F3H activity

The five BnaFLS candidates expressed in seeds were analyzed with respect to conserved amino acids and motifs important for FLS functionality (Figure 4). Both BnaFLS1 candidates contain all conserved amino acids and motifs. All remaining candidates lack the motifs potentially important for FLS activity, namley ‘SxxTxLVP’-, ‘CPQ/RPxLAL’-, and the N-terminal ‘PxxxIRxxxEQP’, in parts or completely. However, all BnaFLS candidates possess the conserved residues for ferrous iron- and 2-oxoglutarate-binding. Only BnaFLS2-1 revealed three amino acid exchanges in the five substrate binding residues analyzed, which are H103N, K173R, and E266D. BnaFLS3-3 and BnaFLS3-4 carry a G235A (G261 in AthFLS1) amino acid exchange. As some FLSs are bifunctional showing F3H-side activity, BnaFLS candidates were additionally screened for residues important for F3H activity (Figure 4). Besides the previously described G235A exchange of both BnaFLS3 candidates, all five BnaFLS candidates possess the residues described to play a role for F3H activity. The high conservation of relevant motifs and amino acids suggested both FLS1 candidates to be bifunctional. Due to the incomplete motifs and exchanges in conserved amino acids of BnaFLS3-3, BnaFLS3-4, and BnaFLS2-1 the FLS and/or F3H activity of these candidates might be affected.

**Figure 4:**
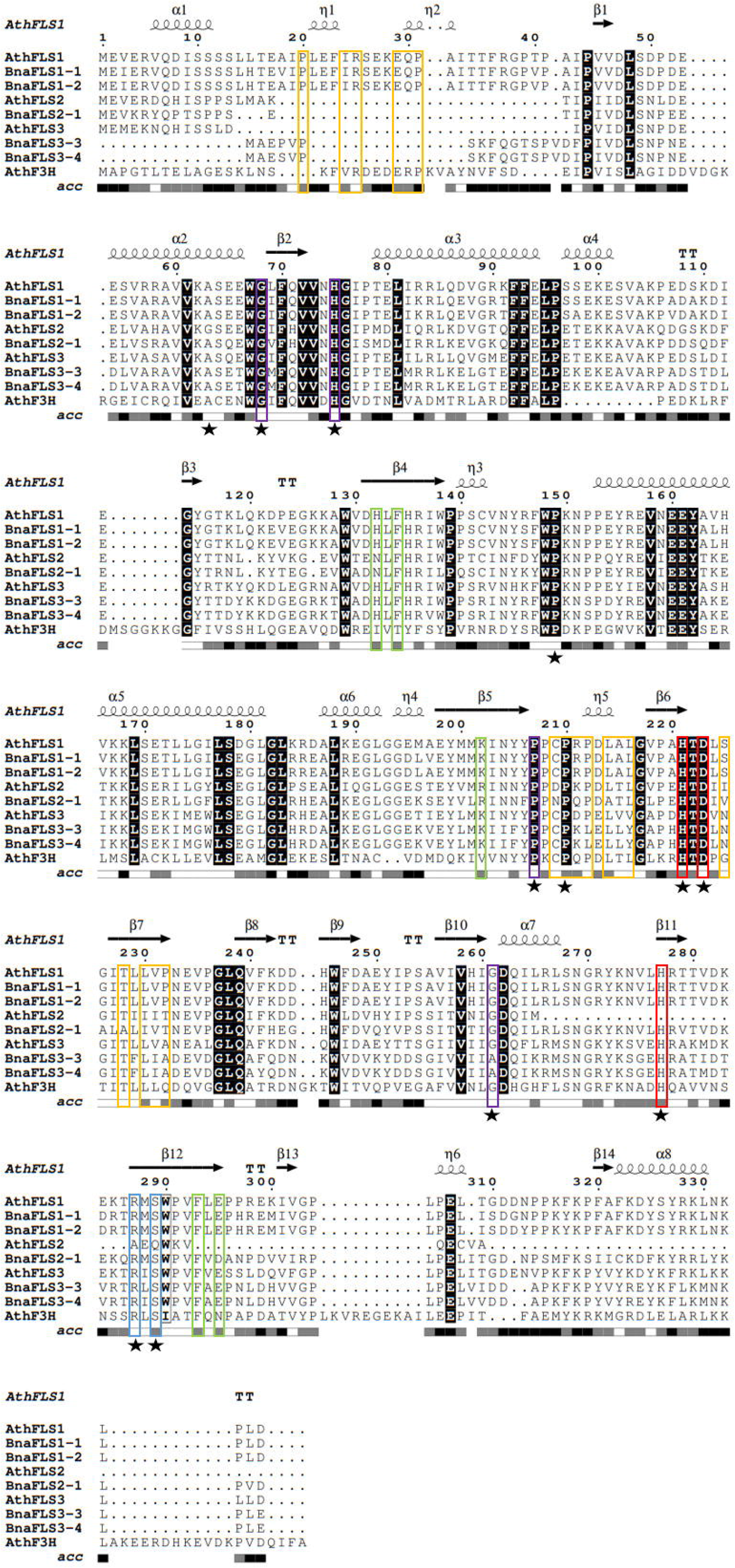
Multiple sequence alignment of BnaFLS candidates relevant for seed flavonol accumulation. Conserved amino acids and motifs important for FLS functionality were labelled as followed: the ‘PxxxIRxxxEQP’, ‘CPQ/RPxLAL’, and ‘SxxTxLVP’ motifs are shown in orange, while residues involved in substrate-, ferrous iron-, and 2-oxoglutarate-binding are marked in green, red, and blue, respectively. Residues important for proper folding and/or highly conserved across 2-ODDs are labelled in violet. Residues relevant for F3H activity are marked with a black star. Black background indicates perfect conservation across all sequences. Secondary structure information is derived from an *in silico* model of AthFLS1 predicted by I-TASSER. acc = relative accessibility.

Moreover, all BnaFLS candidates were predicted to contain no transmembrane helices, signal peptides or N-terminal presequences (mitochondrial-, chloroplast-, thylakoid luminal transfer peptide) and are therefore assumed and predicted to be located in the cytoplasm (Supplementary Table S2, S4, S5).

### 3.6 Functional characterization of BnaFLS candidates

For the functional characterization of BnaFLS1-1, BnaFLS1-2, BnaFLS2-1, BnaFLS3-3, and BnaFLS3-4 *in vivo* bioconversion assays in *E. coli* as well as analysis of stablely transformed *A.thaliana* knock out mutants were performed. The reproducibility of the bioconversion assay was ensured by showing that the observed functionalities of the well-known 2-ODD members AthFLS1, AthFLS3, AthFLS5, AthF3H, and AthANS match literature-based knowledge (Supplementary Figure S6). As expected, AthF3H showed clear F3H activity. In line with previous reports, AthFLS1 was identified as bifunctional possessing FLS activity and F3H side activity and AthANS showed FLS and F3H side activity. None of these activities could be detected for AthFLS5. Although AthFLS3 was reported to have FLS activity under extended assay conditions in *E. coli*, we could not detect FLS or F3H activity.

#### 3.6.1 BnaFLS1-1 and BnaFLS1-2 are bifunctional enzymes exhibiting F3H and FLS activity

The predictions reported above were experimentally validated for BnaFLS1-1 and BnaFLS1-2, which were indeed bifunctional. Both enzymes can generate dihydrokaempferol and kaempferol (Figure 5A-B). To validate bifunctionality *in planta*, flavonol glycosides of the *ans/fls1 A. thaliana* double mutants transgenic for *BnaFLS1-1* and *BnaFLS1-2* were analyzed via HPTLC. In line with the bioconversion assay results, the *in planta* analysis revealed successful complementation of the *ans/fls1 A. thaliana* double knock out mutant by *BnaFLS1-1* or *BnaFLS1-2*, restoring the *A. thaliana* wildtype phenotype (Figure 5C). Additionally, DPBA-staining of young seedlings was used to visualize flavonoid derivatives under UV illumination, including kaempferol (green) and quercetin derivatives (yellow, orange). This *in situ* validation revealed a restoration of the wildtype phenotype by *BnaFLS1-1* and *BnaFLS1-2* compared to the *f3h* and *ans/fls1* knock out mutants (Figure 5D-E). Collectively, these results showed that *BnaFLS1-1* and *BnaFLS1-2* encode bifunctional enzymes, which exhibit FLS and F3H activity.

**Figure 5:**
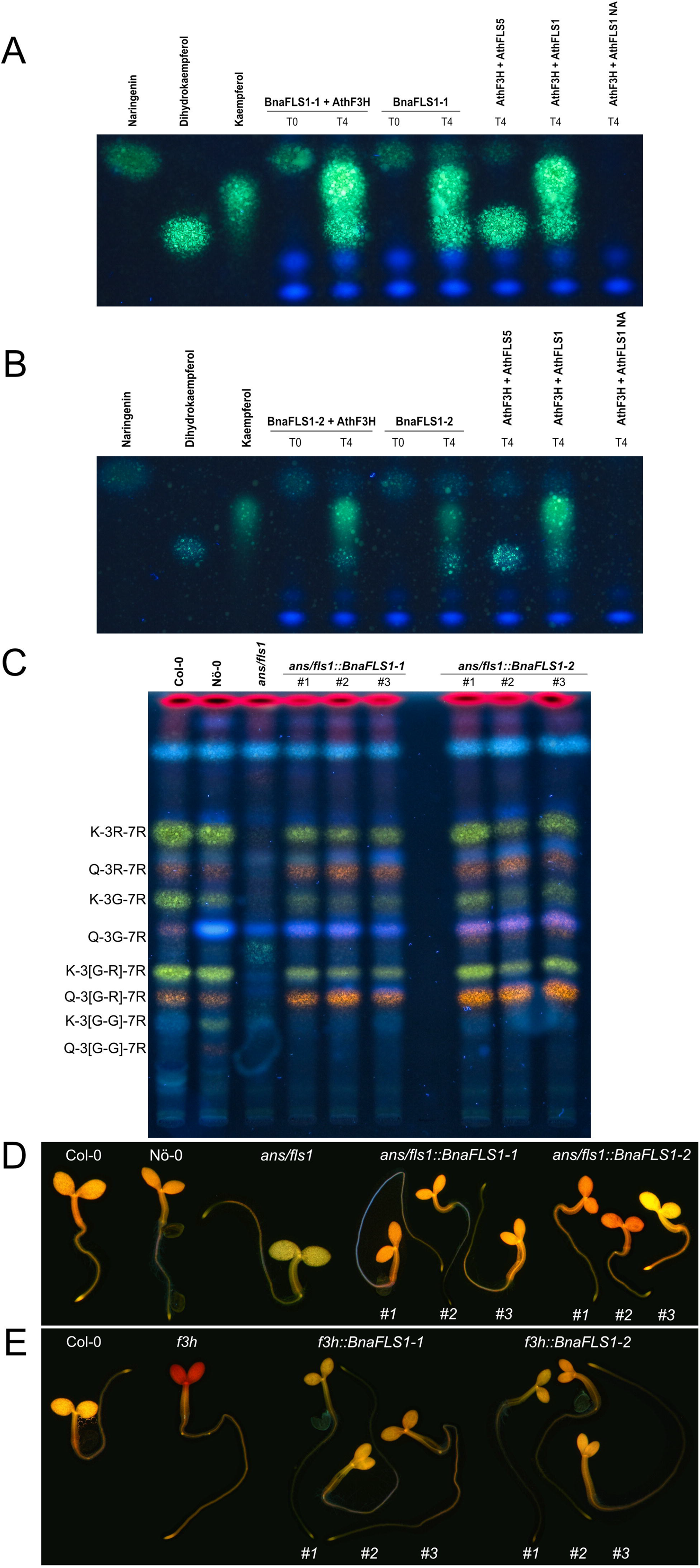
BnaFLS1-1 and BnaFLS1-2 are bifunctional enzymes exhibiting F3H and FLS activity. (A) and (B) Bioconversion assay results based on a HPTLC using extracts from *E. coli* expressing recombinant BnaFLS1-1 or BnaFLS1-2. The substrate of F3H naringenin, as well as the FLS substrate dihydrokaempferol and the product kaempferol were used as standards. AthFLS1 served as positive control and AthFLS5 as negative control. In the last sample no Nargingenin (NA) was supplemented. (C) HPTLC on silica gel-60 plates of methanolic extracts of stem of Col-0, Nö-0, *ans/fls1 A. thaliana* knock out mutant, and three independent T2 *ans/fls1 A. thaliana* knock out *BnaFLS1-1* and *BnaFLS1-2* complementation lines followed by DPBA staining, applied in this order. Pictures were taken under UV illumination. Kaempferol- and quercetin derivatives are green and orange respectively, while sinapate derivates are faint blue, dihydrokaempferol derivates are turquois, and chlorophylls appear red. The following flavonoid derivates are labeled: kaempferol-3-O-rhamnoside-7-O-rhamnoside (K-3R-7R), quercetin-3-O-rhamnoside-7-O-rhamnoside (Q-3R-7R), kaempferol-3-O-glucoside-7-O-rhamnoside (K-3G-7R), quercetin-3-O-glucoside-7-O-rhamnoside (Q-3G-7R), kaempferol-3-O-glucorhamnosid-7-O-rhamnoside (K-3[G-R]-7R), quercetin-3-O-glucorhamnosid-7-O-rhamnoside (Q-3[G-R]-7R), kaempferol-3-O-gentiobioside-7-O-rhamnoside (K-3[G-G]-7R), and quercetin-3-O-gentiobioside-7-O-rhamnoside (Q-3[G-G]-7R). (D) and (E) Flavonol staining in young seedlings of Col-0, Nö-0, *ans/fls1* double and *f3h* single *A. thaliana* knock out mutant, as well as representative pictures of three independent T2 *ans/fls1 A. thaliana* knock out *BnaFLS1-1* and *BnaFLS1-2* complementation lines and three independent T3 *f3h A. thaliana* knock out *BnaFLS1-1* and *BnaFLS1-2* complementation lines. Flavonols in norflurazon-bleached seedlings were stained with DPBA until saturation and imaged by epifluorescence microscopy. Orange color indicates the accumulation of quercetin derivates. Photos of representative seedlings are shown.

#### 3.6.2 BnaFLS family members with divergent enzyme functionalities

Interestingly, only BnaFLS1-1 and BnaFLS1-2 revealed FLS activity out of the five *BnaFLS* candidates expressed in seeds. While neither F3H nor FLS activity could be detected for BnaFLS2-1 (Supplementary Figure S7), both BnaFLS3 candidates showed F3H activity *in vivo* and *in planta*, thus they can convert naringenin to dihydroflavonols (Figure 6A-E). However, no FLS activity could be detected for both BnaFLS3s (Figure 6A-E). These findings validate the predictions based on the presence of almost all important residues for F3H activity for both BnaFLS3s, with G235A (G261 in AthFLS1) being the only exception (Figure 4).

**Figure 6:**
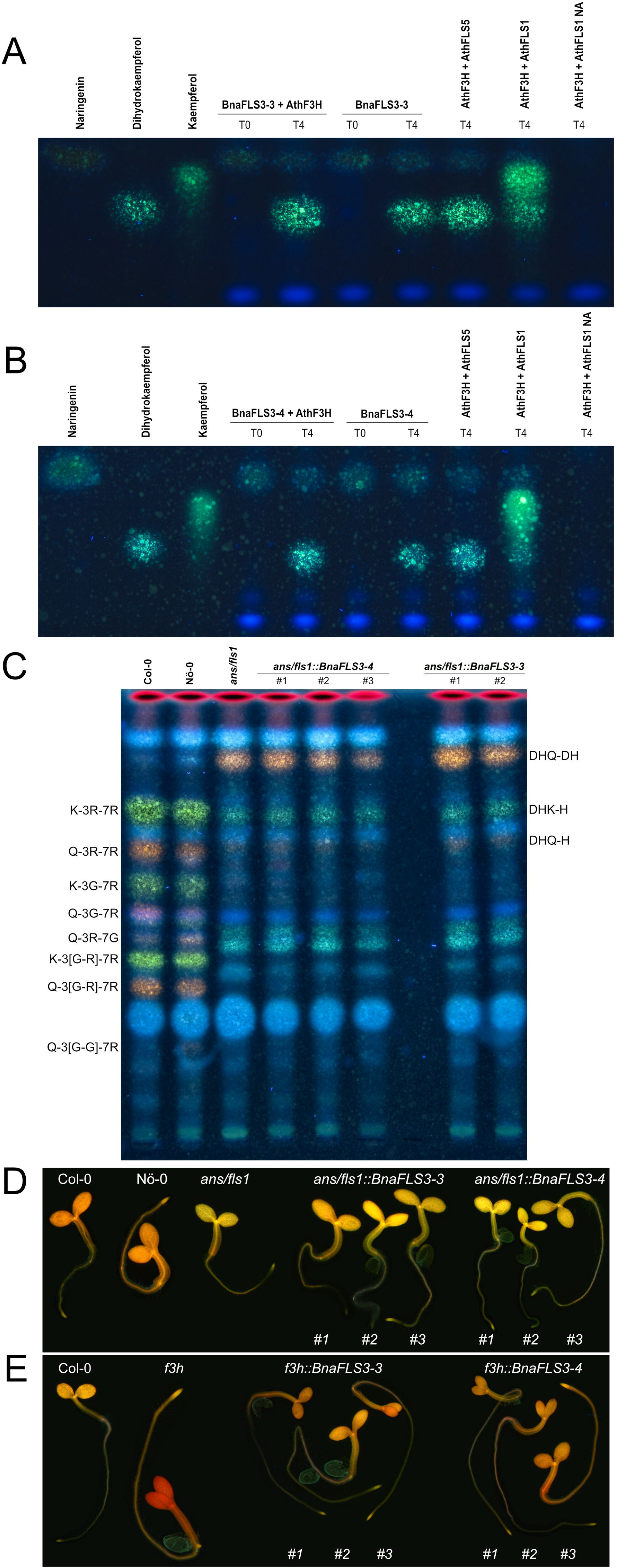
BnaFLS3-3 and BnaFLS3-4 exhibit F3H activity. See Figure 5 for detailed figure description. (A) Bioconversion assay results of BnaFLS3-3 and (B) BnaFLS3-4. (C) The following flavonoid derivates were additionally labeled: dihydroquercetin-deoxyhexoside (DHQ-DH), dihydrokaempferol-hexoside (DHK-H), dihydroquercetin-hexoside (DHQ-H), quercetin-3-O-rhamnoside-7-O-glucoside (Q-3R-7G). (D) and (E) Flavonol staining in young seedlings.

### 3.7 Structural modelling revealed three major differences of the bifunctional enzymes compared to monofunctional ones

To investigate whether the bifunctionality of both BnaFLS1s compared to both BnaFLS3s, which showed only F3H activity, might be based on structural differences *in silico*, 3D models were generated (Figure 7A-F). The BnaFLS1s showed three major differences compared to both BnaFLS3s, which offer insights into the potential mechanisms of bifunctionality: i) Both BnaFLS3 models revealed a shorter N-terminus compared to BnaFLS1s, resulting in the loss of the presumably FLS-specific ‘PxxxIRxxxEQP’-motif and α-helices (Figure 4, Figure 7). ii) The amino acid G261 proposed to be involved in proper folding is only present in both BnaFLS1s, while BnaFLS3s carry an alanine at this position. This residue is located between the transition of a beta-sheet from the jellyroll core structure to an α-helix. The hydrophobic side chain of alanine likely reduces the space in the catalytic center. iii) Both BnaFLS3s show only partial overlaps with the ‘SxxTxLVP’- and ‘CPQ/RPxLAL’-FLS-specific sequence motifs (Figure 4). However, these mismatches do not have a substantial effect on the overall secondary structure in these regions (Figure 7 E-F). Moreover, an extended N-terminus is not essential for F3H activity since it is absent in BnaFLS3-3 and BnaFLS3-4 (Figure 7 D-F).

**Figure 7:**
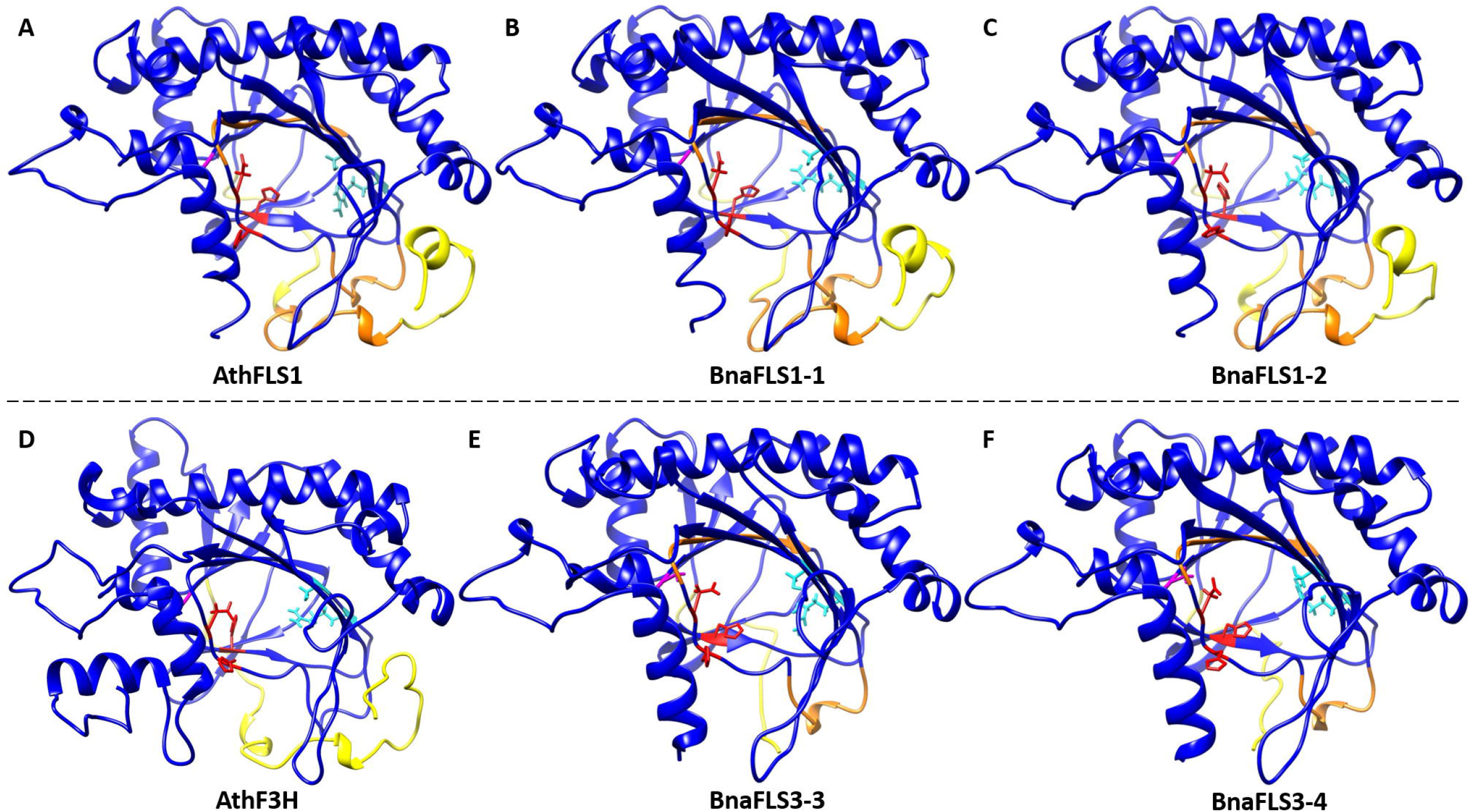
3D secondary structure models of BnaFLS1s and BnaFLS3s. Homology models of (A) AthFLS1, (B) BnaFLS1-1, (C) BnaFLS1-2, (D) AthF3H, (E) BnaFLS3-3, and (F) BnaFLS3-4 modelled via I-TASSER are shown looking into the center of the jellyroll motif. Ferrous iron-coordinating residues are shown in red, 2*-*oxoglutarate binding residues are marked in cyan, and the corresponding position of G261 in AthFLS1 is shown in magenta. The N-terminus divergence between BnaFLS1s and BnaFLS3s is marked in yellow (corresponding to amino acids 1-42 in AthFLS1). Orange regions compromise regions postulated to be specific for FLS.

## 4 Discussion

### 4.1 Phylogeny of *BnaFLS* gene family members

Although flavonols are of agronomical, ornamental, nutritional, and health importance, the major players of the flavonol biosynthesis in the oil and protein crop *B. napus* have not been investigated in great detail yet. So far, only one *FLS* gene was identified via transient expression in tobacco (Vu et al., 2015). However, as there are several members of the *BnaFLS* gene family expressed in seeds, it is necessary to characterize the encoding enzymes to infer which genes contribute to flavonol biosynthesis in *B. napus* seeds.

The members of the *BnaFLS* gene family are more closely related to each other than to any of the other 2-ODDs, which is in line with the results for the *AthFLS* gene family (Owens et al., 2008). In contrast to the *AthFLS* gene family, which is located on chromosome 5 in close proximity (Owens et al., 2008), the members of the *BnaFLS* gene family are distributed across seven chromosomes. Considering the chromosomal rearrangements described for *B. napus* cultivar Darmor-bzh (Chalhoub et al., 2014) and also the chromosomal positions of the *B. rapa* (Guo et al., 2014) and *B. oleracea* (Parkin et al., 2014) *FLS* genes, high local synteny of the *FLS* loci to those of *B. napus* Express 617 was identified. This syntenic relation allowed the assignment of 6 homeologous pairs of the *B. napus FLS* gene family. The homeolog pair *BnaFLS3-3* and *BnaFLS3-4* is located on the pseudochromosome C02 and clusters together with one additional unassigned *BnaFLS3-5* homolog. The position of *BnaFLS3-4* and *BnaFLS3-5* on C02 in the Express 617 assembly likely derives from a mis-assembly as inferred by manual curation of the locus and the frequent assignment of the respective homologs to A02 in other long-read *B. napus* cultivar assemblies like westar and shengli (Song et al., 2020). Moreover, the respective *B. rapa* homologs *BraFLS3-4* (Bra029212) and *BraFLS3-5* (Bra029211) are located on A02. *BraFLS3-4* and *BraFLS3-5* are assumed to originate from duplication of the syntenic *AthFLS2* to *AthFLS5* tandem array. This duplication is part of the whole genome duplication (WGD), but only some genes of the array were retained in *B. rapa* (Guo et al., 2014). *BraFLS3-5* is assumed to be derived from a gene duplication event of *BraFLS3-4* (Guo et al., 2014), which is underlined by the close proximity of the two homologs *BnaFLS3-4* and *BnaFLS3-5* (only 2.9 kbp apart in the Express 617 assembly, see Table 1). Moreover, *BraFLS2-2* (Bra038647) and *BraFLS3-2* (Bra038648) are assumed to have emerged by WGD events as described before (Guo et al., 2014). A similar originating mechanism is assumed for *BolFLS2-1* (Bo3g103270) and *BolFLS3-1* (Bo3g103260) and thus for their respective homologs *BnaFLS3-1* and *BnaFLS2-1*. Therefore, these ancient duplication events shaped the *B. napus FLS* gene family.

In *A. thaliana FLS5* encodes a full-length protein, which contains amino acid exchanges important for hydrogen bonding of the substrate most likely resulting in a non-functional polypeptide (Owens et al., 2008; Preuss et al., 2009). In line with our results, no *FLS5* homolog was identified in *B. rapa* (Guo et al., 2014). However, we could also not detect *FLS5* in Brassiceae, Arabideae, Eutremeae, and Coluteocarpeae, but *FLS5* was detected in the Camelineae, which include *A. thaliana*, as well as in Boechereae. Thus, we postulate that *FLS5* emerged after the divergence of the common ancestor of the parental species of *B. napus* (*B. rapa* and *B. oleracea*) and *A. thaliana* rather than that it was frequently lost after the WGD events of the tandem array as postulated by Guo *et al*. 2014 (Guo et al., 2014).

*FLS6* was characterized as a pseudogene in *A. thaliana* (Owens et al., 2008; Stracke et al., 2009) and no *FLS6* homolog was identified in *B. rapa* (Guo et al., 2014). These findings are in line with our results showing that *FLS6* was lost very recently in *B. rapa* and *B. oleracea* and consequently is not present in *B. napus*, since *FLS6* is still present in *Raphanus sativus*. As *FLS6* was identified as pseudogene and *FLS5* is known to encode a non-functional protein in *A. thaliana* (Owens et al., 2008; Preuss et al., 2009; Stracke et al., 2009), the parental species *B. oleracea* and *B. rapa* have already eliminated these ‘unnecessary’ genes.

However, some *BnaFLS* genes are retained as they still encode functional proteins like *BnaFLS3-3* and *BnaFLS3-4*, which encode for proteins with F3H activity. Importantly, both BnaFLS3s show a higher sequence identity with functional FLSs compared to F3H homologs, although exhibiting only F3H activity. This fact provides clear evidence that a classification solely based on amino acid sequences is not sufficient to infer functionalities of FLS family members and very likely 2-ODDs in general.

### 4.2 The *BnaFLS* gene family contains two bifunctional FLSs

Bifunctionality has so far not been reported for a FLS from *B. napus*. By using two independent methods, we demonstrated bifunctionality of the two BnaFLS1 homeologs, which exhibit F3H and FLS activity. Thus, BnaFLS1-1 and BnaFLS1-2 are responsible for flavonol production *in planta*. We hypothesize that the respective orthologs of *B. oleracea* (Bo9g174290) and *B. rapa* (Bra009358) are bifunctional enzymes as well (Supplementary Table S6). Moreover, two additional members of the *BnaFLS* gene family have been functionally characterized. Interestingly, BnaFLS3-3 and BnaFLS3-4 revealed only F3H activity, while no FLS activity was detected. By incorporating sequence and structural analyses of 3D secondary structure models of BnaFLS1s vs BnaFLS3s, we proposed a set of evolutionary events underlying the mechanisms of bifunctionality. Both BnaFLS3s lack several amino acids at the beginning of the N-terminus, which could cause the loss of FLS activity as it harbours the ‘PxxxIRxxxEQP’ motif. This motif was previously proposed to be important for FLS activity as it distinguishes FLS from 2-oxoglutarate-/FeII-dependent dioxygenases with other substrate specificities (Owens et al., 2008; Stracke et al., 2009). Additional support for the relevance of this N-terminal region is provided by an AthFLS1 protein lacking the first 21 amino acids which showed no FLS activity (Pelletier et al., 1999; Owens et al., 2008). Moreover, amino acid exchanges in the ‘CPQ/RPxLAL’- and ‘SxxTxLVP’-motif in both BnaFLS3s possibly impact FLS activity. In addition, both BnaFLS3s carry a G235A (G261 in AthFLS1) amino acid exchange in comparison to BnaFLS1s, which might be relevant for bifunctionality as this exchange reduced the activity of a mutated *Citrus unshiu* FLS by 90% (Wellmann et al., 2002). This glycine is conserved across 2-ODDs and is suggested to play a role in proper folding (Wellmann et al., 2002). In accordance, we identified the A235 of BnaFLS3s and G261 of BnaFLS1 located between the transition of a beta-sheet from the jellyroll core structure to an α-helix, thereby the hydrophobic side chain of the alanine might reduce the space in the catalytic center. We propose that FLS bifunctionality is likely influenced by a combination of the identified motifs and residues rather than a single causative change as observed before for other flavonoid enzymes (Gebhardt et al., 2007; Seitz et al., 2007). The impact of each motif or amino acid on FLS bifunctionality needs further investigations that go beyond this study. As these sequence differences of BnaFLS3s do not abolish F3H activity, we uncovered that a truncated N-terminus and G261 are not essential for F3H activity. This is of importance as G261 was reported to be important for F3H activity (Britsch et al., 1993) while it may only play a minor role in conservation of F3H activity.

In addition to the FLS activity of BnaFLS1s, the 2-ODD member ANS might be able to contribute to flavonol production, as AthANS exhibit FLS and F3H side activities *in vitro* (Turnbull et al., 2004). *In planta*, FLS is the major enzyme in flavonol production as AthANS was not able to fully substitute AthFLS1 *in vivo* which is visible in the flavonol deficient *fls1-2* mutant (Owens et al., 2008; Stracke et al., 2009).

*BnaFLS2-1* is most likely a pseudogene. Although *BnaFLS2-1* is still marginally expressed as shown by RNA-Seq data, it carries amino acid exchanges within 3/5 substrate binding residues in addition to a truncated N-terminus, which render the protein non-functional. In addition, an alternative transcript of *BnaFLS2-1* was discovered (Supplementary Figure S5) that leads to a frameshift and thus likely encodes a non-functional protein as well. In *A. thaliana*, a heterologous expressed mutated FLS carrying one of the identified amino acid exchanges, namely K202R (K173R in BnaFLS2-1) is described to possess only 12% of the wild type FLS activity (Chua et al., 2008). In accordance, *AthFLS2* encodes a most likely non-functional protein, which also harbors a truncated N-terminus (Owens et al., 2008). We assume that *BnaFLS2-1* might be derived from a gene duplication event, losing its original function over time due to a pseudogenisation process similar to that proposed for the *AthFLS* gene family members (Preuss et al., 2009; Stracke et al., 2009). The rather low expression of *BnaFLS2-1* across various organs supports this hypothesis.

### 4.3 *BnaFLS1s* are major players in flavonol biosynthesis in *B. napus* seeds

The spatio-temporal patterns of flavonol accumulation in *B. napus* are characterized by the activity of multiple *BnaFLS* genes. Both *BnaFLS3s* are expressed in early seed development while *BnaFLS1s* are expressed during late seed development (Table 2). The similar expression patterns of both *BnaFLS1s* are expected because they are homeologs. Thus, their expression patterns in the parental species *B. rapa* and *B. oleraceae* were likely to be very similar as they fulfill similar functions. In line with these results, *BnaFLS1-1* and *BnaFLS1-2* share co-expressed genes of the flavonoid and phenylpropanoid pathway. Both *BnaFLS1s* are co-expressed with *MYB111*, a regulator of flavonol biosynthesis (Stracke et al., 2007) and contain SG7 MRE in their putative promoter regions. Additionally, genes important for flavonoid transport into the vacuole and anthocyanidin/flavonol glycosylation like *GSTF12* (TT19) and *UGT84A2* (Kitamura et al., 2004; Yonekura-Sakakibara et al., 2012) were identified to be co-expressed with *BnaFLS1s*. These results further support the role of BnaFLS1-1 and BnaFLS1-2 as major players of flavonol biosynthesis in *B. napus* seeds. Moreover, transcriptomic and functional analysis of *BnaFLS1s* indicate gene redundancy.

Both *BnaFLS1s* were mainly expressed in reproductive organs as observed for *AthFLS1* (Owens et al., 2008). *BnaFLS3-4* was identified to be co-expressed with the well-known transcription factors MYB61, MYB123, and MYB5 which play a role in flavonoid biosynthesis and seed coat development in *A. thaliana* (Penfield et al., 2001; Li et al., 2009; Xu et al., 2014). This indicates a likely conserved transcriptional regulation between these two closely related species and supports the importance of flavonols during reproductive processes, e.g. pollen tube growth (Muhlemann et al., 2018).

In line with metabolomic studies showing that phenolic and flavonoid seed content maximized 35 days after flowering (DAF) (Wang et al., 2018), the expression of *BnaFLS1s* was higher at 35 DAF compared to 23 DAF. In accordance, most kaempferol and quercetin derivates reach their abundance peak at 35 DAF (Wang et al., 2018). Thus the expression pattern of *BnaFLS1s* fit well with the flavonol accumulation pattern of developing seeds, where flavonols contribute to seed quality (Wang et al., 2018).

Finally, the expression of *BnaFLSs* family members is not restricted to seeds. Some *BnaFLSs* were identified to be expressed in roots including *BnaFLS3-3* and *BnaFLS3-4* indicating a role of those *BnaFLS* family members in flavonoid biosynthesis in roots.

### 4.4 Future perspectives in engineering flavonol content in *B. napus*

Engineering and breeding of flavonol content is of agronomical, economical, and ornamental importance (Takahashi et al., 2007; Cook et al., 2013; Yin et al., 2019). Flavonol content in petals influences pollinator attraction and drives microevolution of pollinators (Sheehan et al., 2015; Grotewold, 2016). Moreover, flavonols possess ROS scavenging activities and provide protection against UV-B radiation (Harborne and Williams, 2000). Besides the potential of engineering flavonol biosynthesis, anthocyanin and proanthocyanindin production can be engineered as FLS and DFR compete for substrates (Figure 8), thereby influencing important agronomical traits e.g. seed color (Luo et al., 2016). This study identified two bifunctional BnaFLS1s which are highly expressed in seeds and can thus be harnessed to engineer the metabolic flux of seed flavonol biosynthesis in the future (Figure 8). For example, the main bitter off-taste component in rapeseed protein isolates is kaempferol 3-O-(2□-O-Sinapoyl-β-sophoroside) (Hald et al., 2019). Thus, the results of this study provide the basis for breeding low-phenolics lines with focus on the reduction of e.g. kaempferols in seeds, thereby supporting the use of rapeseed protein in human consumption.

**Figure 8:**
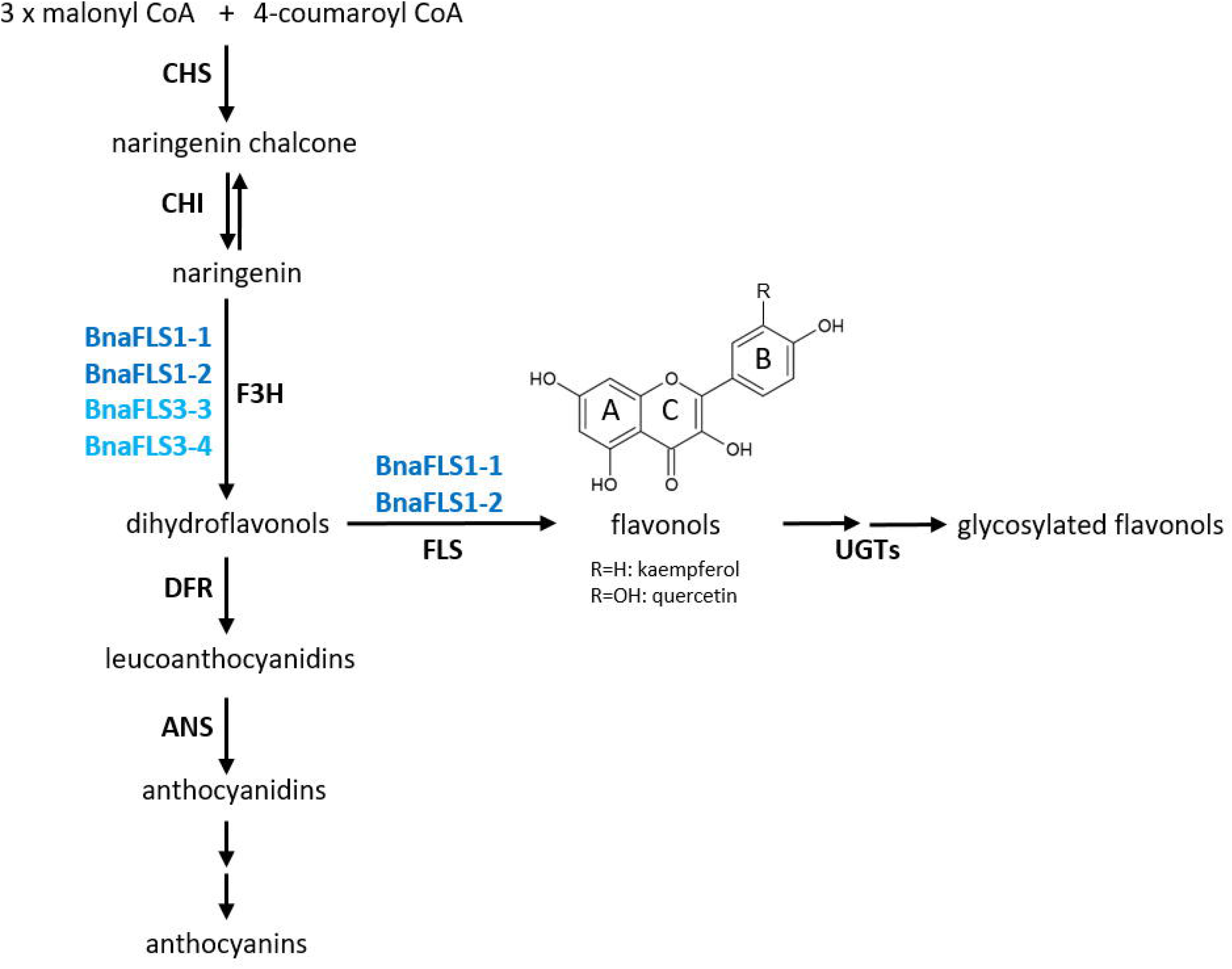
Functional activities of the *B. napus* flavonol synthase family. BnaFLS1-1 and BnaFLS1-2 marked in dark blue, are bifunctional enzyme exhibiting F3H and FLS activity. BnaFLS3-3 and BnaFLS3-4 labelled in light blue possess F3H activity.

## 5 Data Availability Statement

The RNA-Seq data sets generated for this study can be found in the ENA/NCBI BioProject PRJEB45399.

## 7 Author Contributions

HMS, DH and BW conceived and designed research. HMS, MS, TB, MB and PV investigated and conducted experiments. HMS performed bioinformatic analyses and data curation. HMS wrote the initial draft manuscript. HMS, BW, DH, MB, and TB revised the manuscript.

## 9 Acknowledgments

We are extremely grateful to all researchers who submitted their *B. napus* RNA-Seq data sets to the appropriate databases and published their experimental findings. We thank Ralf Stracke for critical proof-reading and discussion. Moreover, we are grateful to Andrea Voigt for excellent technical assistance. In addition, we thank Nele Tiemann for her help in constructing *BnaFLS1-2* plasmids. We thank Rod Snowdon and Huey Tyng Lee for their support and early excess to the *B. napus* Express 617 reference genome sequence. We thank Christian Möllers for supporting us with viable seeds of Express 617. We thank the Sequencing Core Facility for doing an excellent job in determining DNA sequences. We thank the Center for Biotechnology (CeBiTec) at Bielefeld University for providing an environment to perform the computational analyses.

## 10 Supplementary Material

**Supplementary Table S1: Brassicaceae data sets used for phylogenetic analysis**.

**Supplementary Table S2: Gene-specific features of *BnaFLS* genes**.

**Supplementary Table S3: Theoretical isoelectric point and molecular weight of analyzed proteins**.

**Supplementary Table S4: TargetP-2.0 prediction results**.

**Supplementary Table S5: SignalP-5.0 prediction results**.

**Supplementary Table S6: Amino acid sequence identities of BnaFLSs compared to FLS orthologs of *A. thaliana, B. rapa*, and *B. oleracea***.

**Supplementary Table S7: SRA data sets used for organ-specific RNA-Seq analysis**.

**Supplementary Table S8: SRA data sets used for condition-independent co-expression analysis**.

**Supplementary Table S9: Functional annotation used in this work**.

**Supplementary Table S10: Oligonucleotide primers used in this work**.

**Supplementary Table S11: Genes co-expressed with *BnaFLS1-1***.

**Supplementary Table S12: Genes co-expressed with *BnaFLS1-2***.

**Supplementary Table S13: Genes co-expressed with *BnaFLS2-1***.

**Supplementary Table S14: Genes co-expressed with *BnaFLS3-3***.

**Supplementary Table S15: Genes co-expressed with *BnaFLS3-4***.

**Supplementary Figure S1: Phylogeny of BnaFLS candidates and other plant 2-ODDs**.

Relative bootstrap-values are shown next to relevant nodes. The phylogenetic tree is based on amino acid sequences of the 2-ODD members FLS, ANS, F3H, and 2-ODD-like sequences derived from Kawai *et al*. 2014.

**Supplementary Figure S2: MYB recognition elements of *BnaFLSs***.

MYB recognition elements upstream of the transcriptional start site (black arrow) annotated based on newly generated RNA-Seq data of *BnaFLS* genes are shown. The SG7 consensus MRE (5’-AcCTACCa-3’/5’-tGGTAGgT-3’) is marked in blue, while the MRE of MYB24 (5’-CNGTTR-3’/5’-RAACNG-3’) is shown in green. Bases differentiating from the consensus motif are underlined. The start codon is highlighted in yellow.

**Supplementary Figure S3: SDS-PAGE of recombinant proteins analyzed in this work**. Recombinant proteins are marked with a red arrow. The *E. coli* strain BL21 was used as control. uninduced (-), induced (+).

**Supplementary Figure S4: Genomic structure of *BnaFLSs***.

The exon-intron structure of *BnaFLSs* is shown. The exons are split into coding sequences (CDS, black) and untranslated regions (UTR, gray) and are displayed by rectangles, introns are displayed as black connecting lines.

**Supplementary Figure S5: Genomic structure of the alternative transcript of *BnaFLS2-1.2***.

(A) The exon-intron structure of *BnaFLS2-1.1 and BnaFLS2-1.2* is shown. The coding sequences (CDS, black), untranslated regions (UTR, gray), and introns are displayed by black and gray rectangles, as well as black connecting lines, respectively. The alternative transcript *BnaFLS2-1.2* contains an additional third exon of 173 bp rendering the encoded 253 amino acid protein most likely non-functional. (B) A frameshift causes a nonsense mutation in the additional third exon. This transcript was observed in seed samples (23 DAF and 35 DAF).

**Supplementary Figure S6: Bioconversion assays of *A. thaliana* 2-ODD members**.

(A) and (B) Bioconversion assay results based on a HPTLC using extracts from *E. coli* expressing recombinant AthFLS1 or AthFLS3, respectively. The substrate of F3H naringenin, as well as the FLS substrate dihydrokaempferol and the product kaempferol were used as standards. AthFLS1 served as positive control and AthFLS5 as negative control. In the last sample no Nargingenin (NA) was supplemented. (C) Bioconversion assay results of AthF3H and AthFLS5. The *E. coli* strain BL21 was used as control.

**Supplementary Figure S7: Functional characterization of BnaFLS2-1**.

See Figure 5 for detailed figure description. (A) Bioconversion assay results of BnaFLS2-1. (B) The following flavonoid derivates were additionally labeled: dihydroquercetin-deoxyhexoside (DHQ-DH), dihydrokaempferol-hexoside (DHK-H), dihydroquercetin-hexoside (DHQ-H), quercetin-3-O-rhamnoside-7-O-glucoside (Q-3R-7G). (C) and (D) Flavonol staining in young seedlings.

**Supplementary File S1: List of plant 2-ODDs amino acid sequences used in phylogenetic analysis**.

**Supplementary File S2: 3D secondary structure models used in this work**.

**Supplementary File S3: Corrected structural annotation of *BnaFLS* genes**.

**Supplementary File S4: CDS of BnaFLSs from this work**.

## 11 Contribution to the field statement

Rapeseed is the second most important oil crop worldwide. The presence of anti-nutritional components renders rapeseed protein, which remains after oil extraction, unusable for human consumption. Flavonols are a major group of phenolics and contribute to the anti-nutritional components. Flavonol biosynthesis branches of from flavonoid biosynthesis. Previous studies in *B. napus* were mainly focused on metabolites, or cover analyses of enzymes/genes action in early steps of flavonoid biosynthesis, preceding flavonol synthase (FLS). In this work, we identified the members of the rapeseed *FLS* gene family and discuss the underlying evolutionary events that shaped the *FLS* gene family. The *FLS* gene family members were analyzed at genomic and transcriptomic level. As seeds are the major organ of agronomical importance, we focussed on *FLS* genes expressed in seeds. These candidates were functionally characterized using *in vivo* and *in planta* experiments. BnaFLS1-1 and BnaFLS1-2 were identified as bifunctional enzymes exhibiting FLS- and F3H activity. Potential mechanisms underlying bifunctionality are presented. Finally, the findings are discussed in the light of the flavonol biosynthesis in *B. napus* pointing towards future directions e.g. to support the use of rapeseed protein in human consumption.

