## Supplementary figures and images for "Characterization of the *Brassica napus* flavonol synthase gene family reveals bifunctional flavonol synthases"

### Supplementary Figure S1

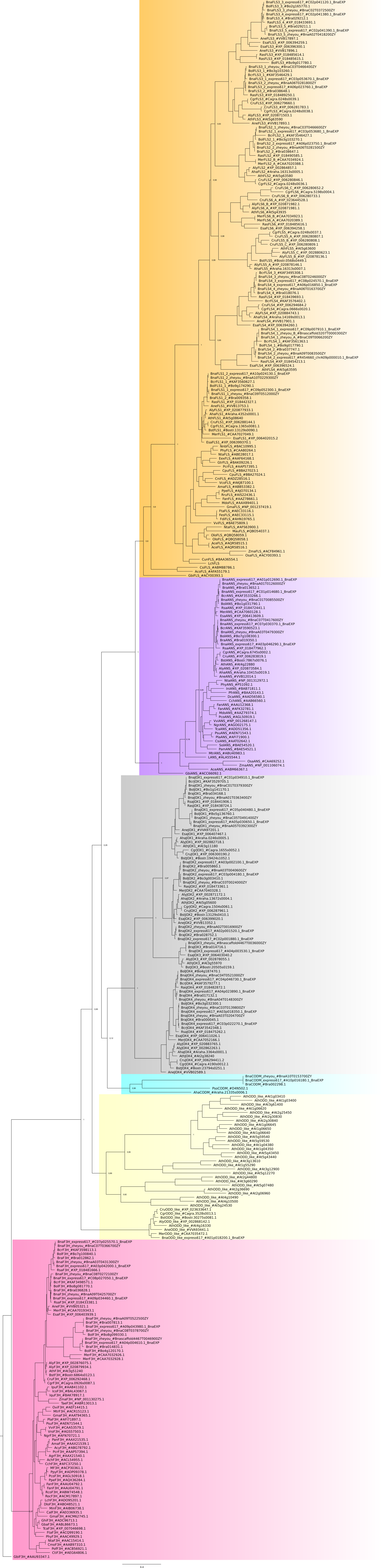

### Supplementary Figure S2

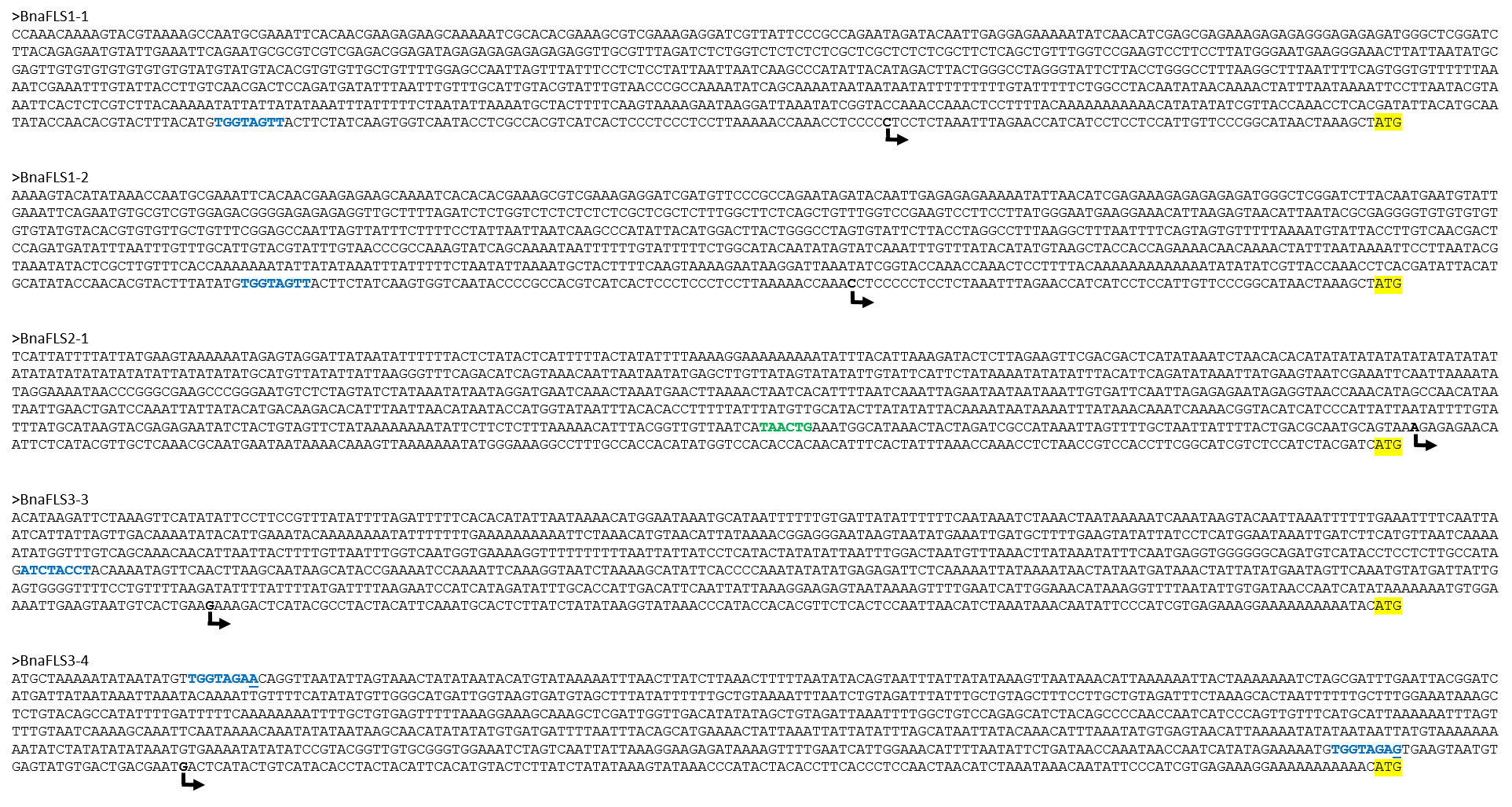

### Supplementary Figure S3

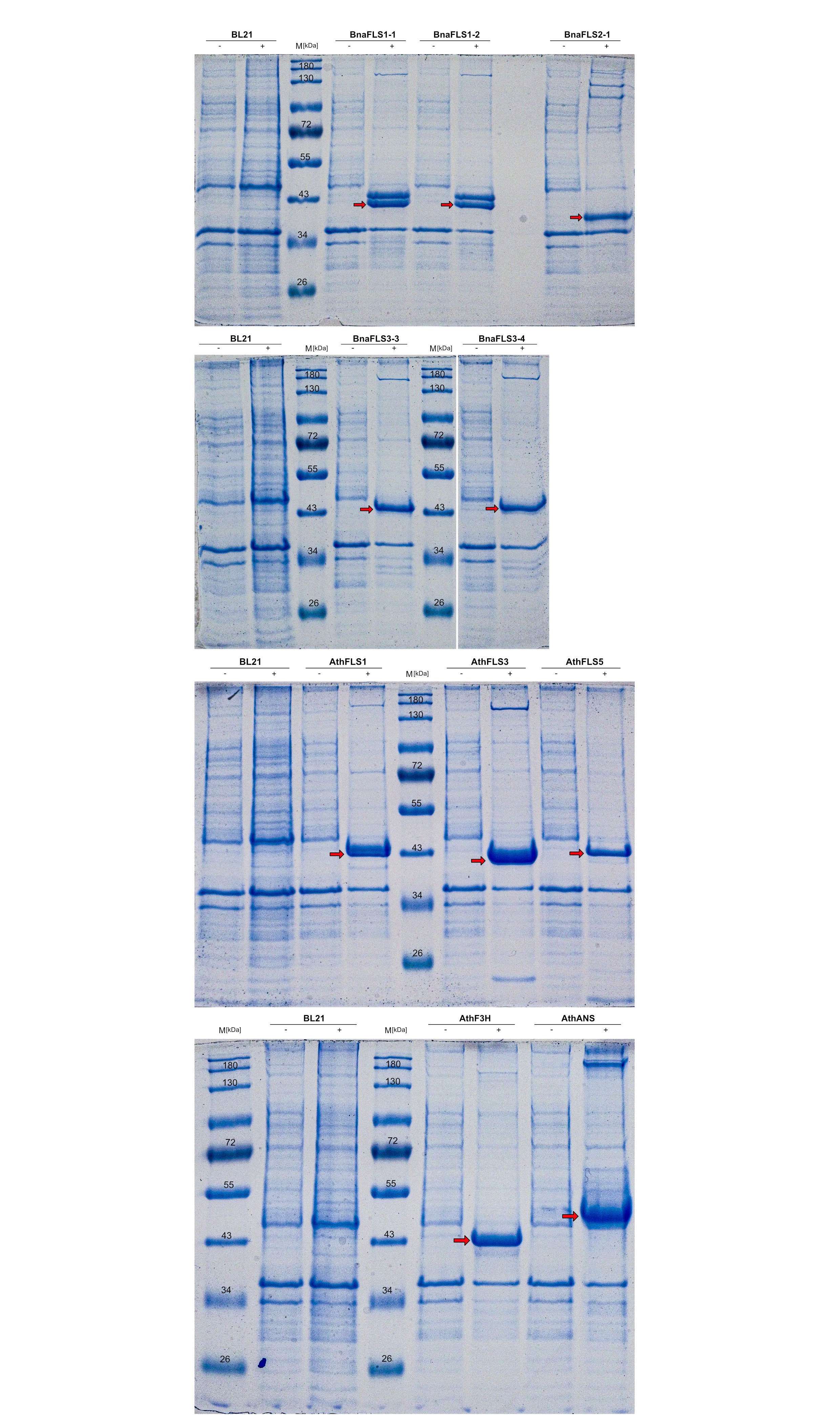

### Supplementary Figure S4

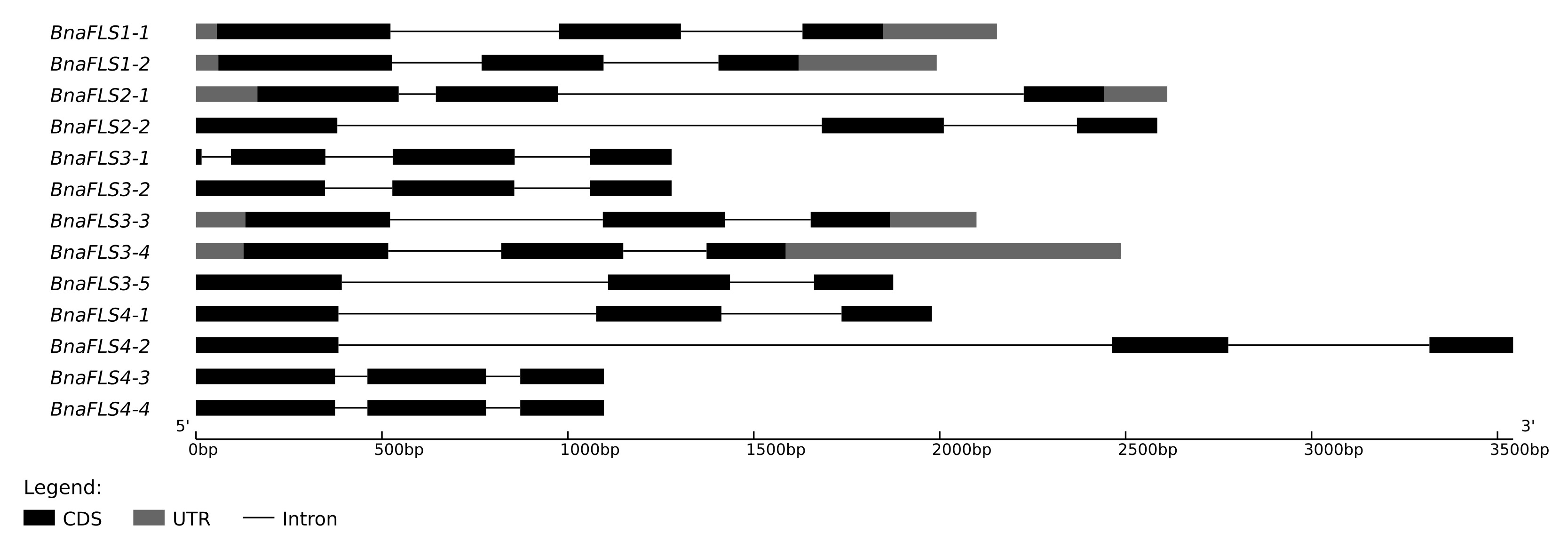

### Supplementary Figure S5

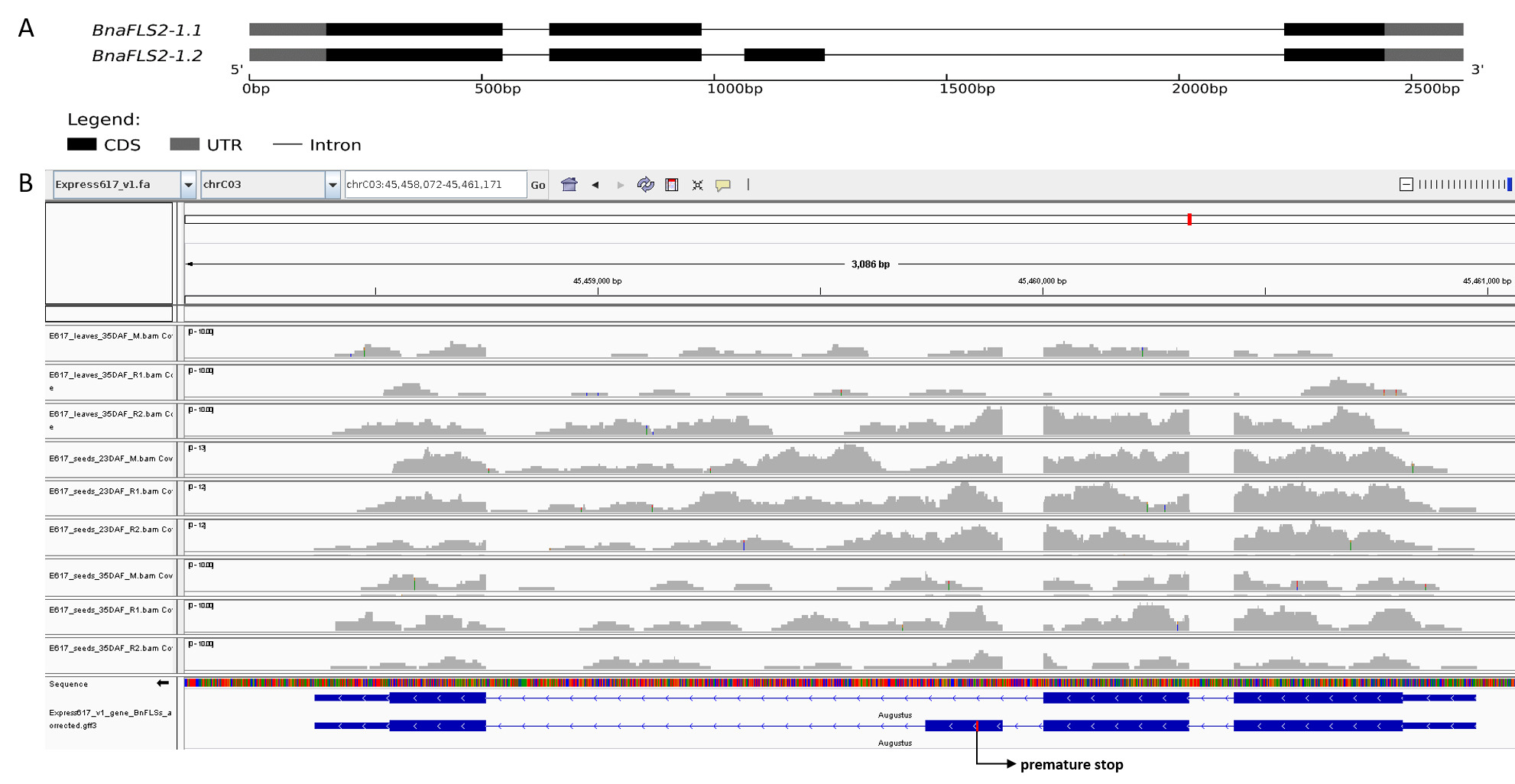

### Supplementary Figure S6

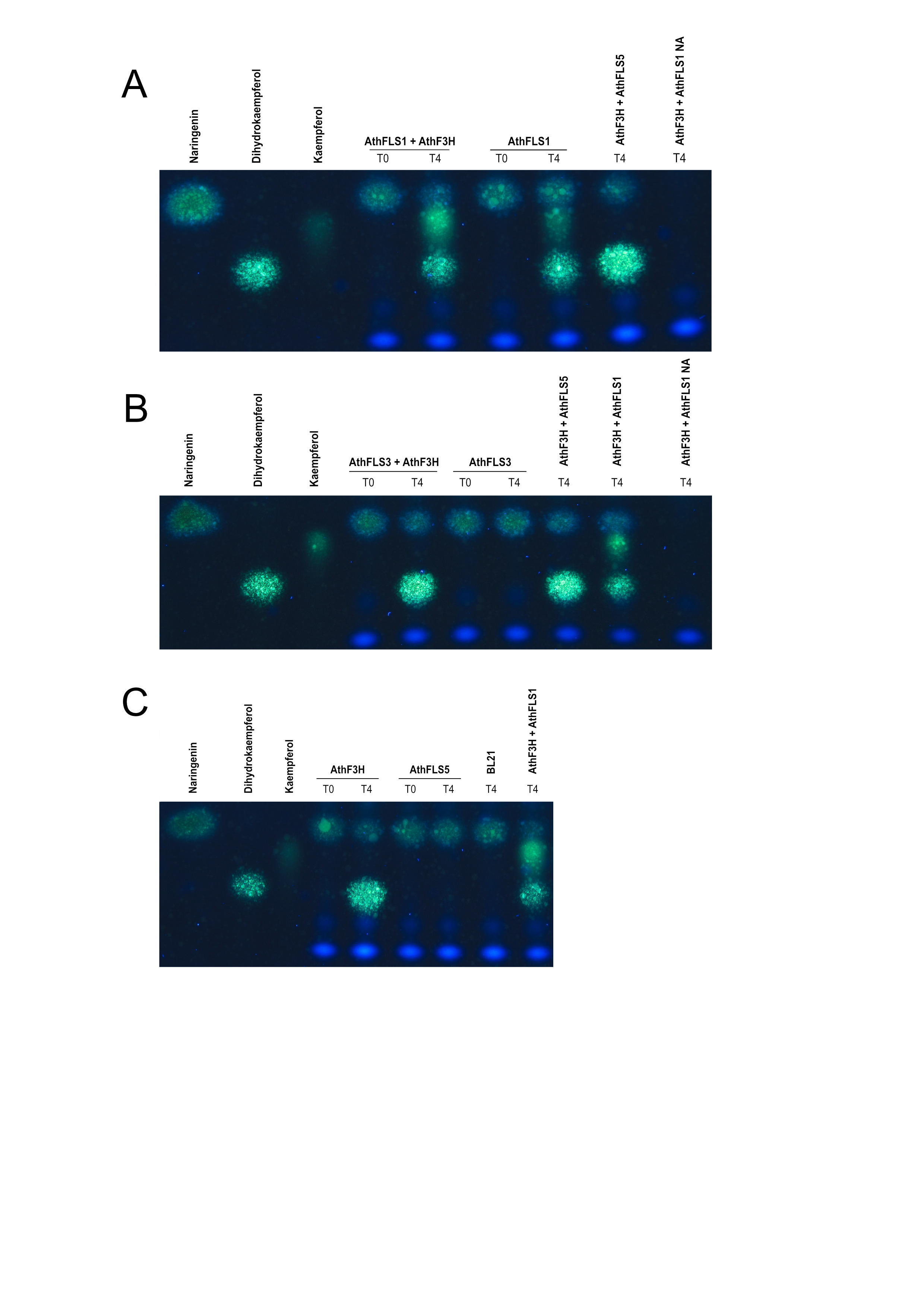

### Supplementary Figure S7

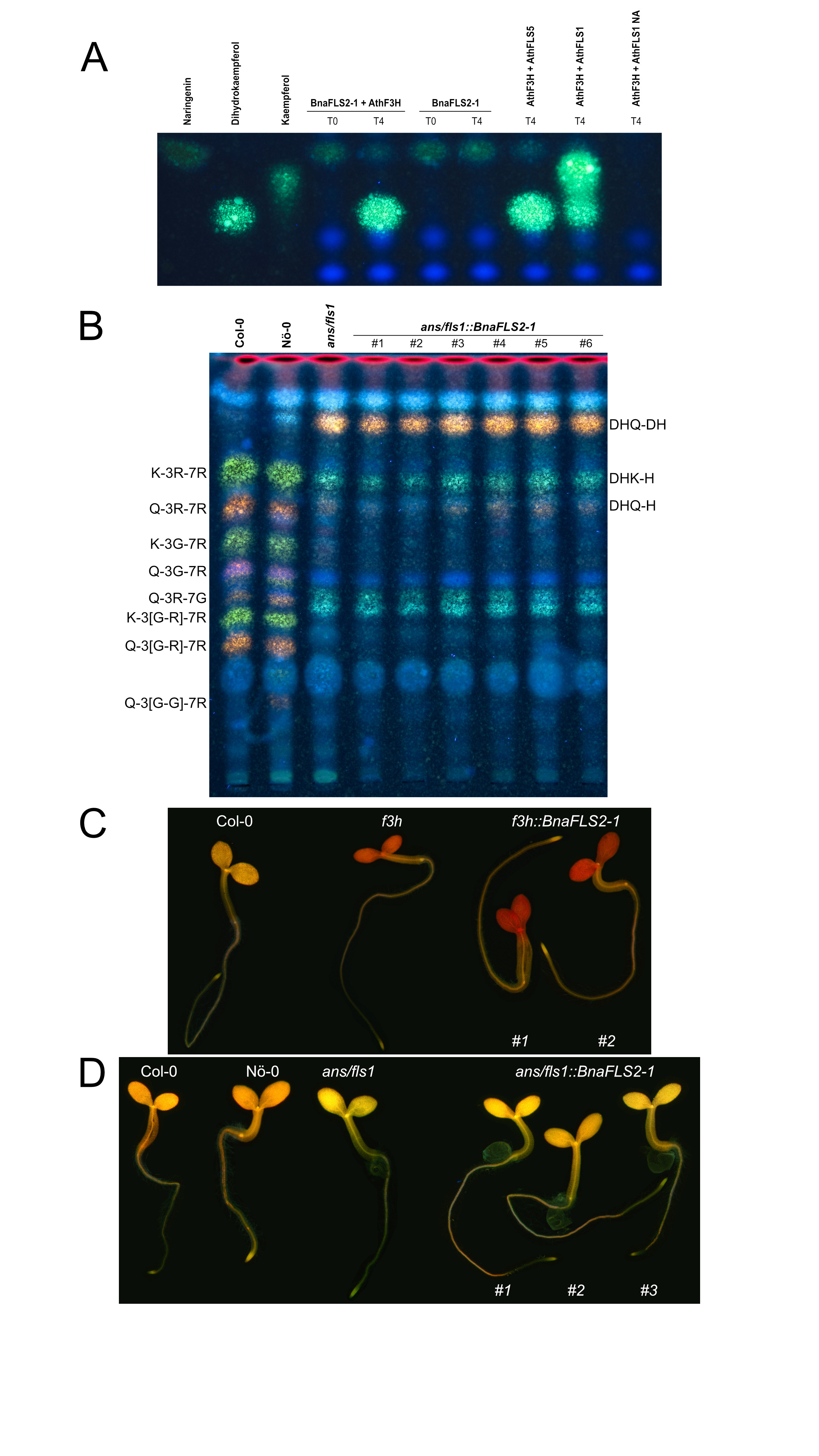
